# Divergent Proteome Reactivity Influences Arm-Selective Activation of Pharmacological Endoplasmic Reticulum Proteostasis Regulators

**DOI:** 10.1101/2023.01.16.524237

**Authors:** Gabriel M. Kline, Ryan J Paxman, Chung-Yon Lin, Nicole Madrazo, Julia M. Grandjean, Kyunga Lee, Karina Nugroho, Evan T. Powers, R. Luke Wiseman, Jeffery W. Kelly

**Author notes:** These authors contributed equally (Gabriel M. Kline, Ryan J Paxman). To whom correspondence should be addressed: Jeffery W. Kelly Department of Chemistry The Scripps Research Institute; R. Luke Wiseman Department of Molecular Medicine The Scripps Research Institute La Jolla, CA 92037.

## Abstract

Pharmacological activation of the activating transcription factor 6 (ATF6) arm of the Unfolded Protein Response (UPR) has proven useful for ameliorating proteostasis deficiencies in a variety of etiologically diverse diseases. Previous high-throughput screening efforts identified the small molecule AA147 as a potent and selective ATF6 activating compound that operates through a mechanism involving metabolic activation of its 2-amino-*p*-cresol substructure affording a quinone methide, which then covalently modifies a subset of ER protein disulfide isomerases (PDIs). Intriguingly, another compound identified in this screen, AA132, also contains a 2-amino-*p*-cresol moiety; however, this compound showed less transcriptional selectivity, instead globally activating all three arms of the UPR. Here, we show that AA132 activates global UPR signaling through a mechanism analogous to that of AA147, involving metabolic activation and covalent PDI modification. Chemoproteomic-enabled analyses show that AA132 covalently modifies PDIs to a greater extent than AA147. Paradoxically, activated AA132 reacts slower with PDIs, indicating it is less reactive than activated AA147. This suggests that the higher labeling of PDIs observed with activated AA132 can be attributed to its lower reactivity, which allows this activated compound to persist longer in the cellular environment prior to quenching by endogenous nucleophiles. Collectively, these results suggest that AA132 globally activates the UPR through increased engagement of ER PDIs. Consistent with this, reducing the cellular concentration of AA132 decreases PDI modifications and allows for selective ATF6 activation. Our results highlight the relationship between metabolically activatable-electrophile stability, ER proteome reactivity, and the transcriptional response observed with the enaminone chemotype of ER proteostasis regulators, enabling continued development of next-generation ATF6 activating compounds.

## INTRODUCTION

The unfolded protein response (UPR) is an endoplasmic reticulum (ER) stress-responsive signaling pathway that corrects imbalances in ER protein homeostasis (proteostasis) caused by an ER stress.^1–3^ The UPR functions by simultaneously expanding ER folding capacity and restricting ER protein flux to restore ER proteostasis through both transcriptional and non-transcriptional responses.^1, 3, 4^ Activation of the UPR occurs downstream of three resident ER stress sensors: PKR-like Endoplasmic Reticulum Kinase (PERK), Inositol-requiring enzyme 1 (IRE1), and Activating Transcription Factor 6 (ATF6). Transcription factors regulated downstream of ATF6 and IRE1, ATF6f and XBP1s respectively, induce expression of numerous protective genes that remodel biological pathways involved in cellular metabolism, redox regulation, and ER proteostasis.^1, 3, 4^ This regulation functions to alleviate the ER stress and adapt cellular physiology to pathologic ER insults. While persistent activation of the IRE1 and PERK arms of the UPR have been associated with pathologic consequences, constitutive ATF6 activation to physiologically relevant levels has not generally been found to be detrimental in mammalian cell culture or mouse models.^5–10^ As such, genetic and pharmacological activation of ATF6 signaling has proven beneficial in mitigating pathological conditions resulting from proteostasis imbalances in numerous disease models.^11–16^ This suggests that ATF6 is an attractive therapeutic target to intervene in etiologically diverse diseases.^4, 5, 17, 18^

We previously conducted a cell-based high-throughput screen to identify selective activators of ATF6 signaling.^19^ From this screen, N-(2-hydroxy-5-methylphenyl)-3-phenylpropanamide (AA147; **Fig 1A**) emerged as a selective ATF6 activator that is able to promote adaptive ATF6 activity to mitigate pathologies associated with etiologically diverse disorders.^11, 19–26^ We found that AA147 functions as a prodrug, wherein oxidative conversion of the 2-amino-*p*-cresol substructure by ER-resident oxidases (e.g., cytochrome P450s) leads to an electrophilic quinone methide and selective covalent engagement of a subset of ER proteins primarily consisting of protein disulfide isomerases (PDIs).^27^ PDIs maintain ATF6 in disulfide-bonded structures that limit its activation.^28–31^ This suggested that AA147-dependent modification of a subset of PDIs could lead to reduction, monomerization, and subsequent trafficking of ATF6 to the Golgi, where S1/S2-enabled proteolytic cleavage affords the cytosolic ATF6 transcription factor amenable to nuclear localization and transcriptional remodeling. Consistent with this, we showed that genetic manipulation of PDIs impaired AA147-dependent ATF6 activation, clearly linking compound modifications of PDIs to the selective ATF6 transcriptional activity observed for this compound.^27^ However, the chemical properties of AA147 allowing PDI modification and subsequent selective ATF6 activation remain to fully established.

**Figure 1.**
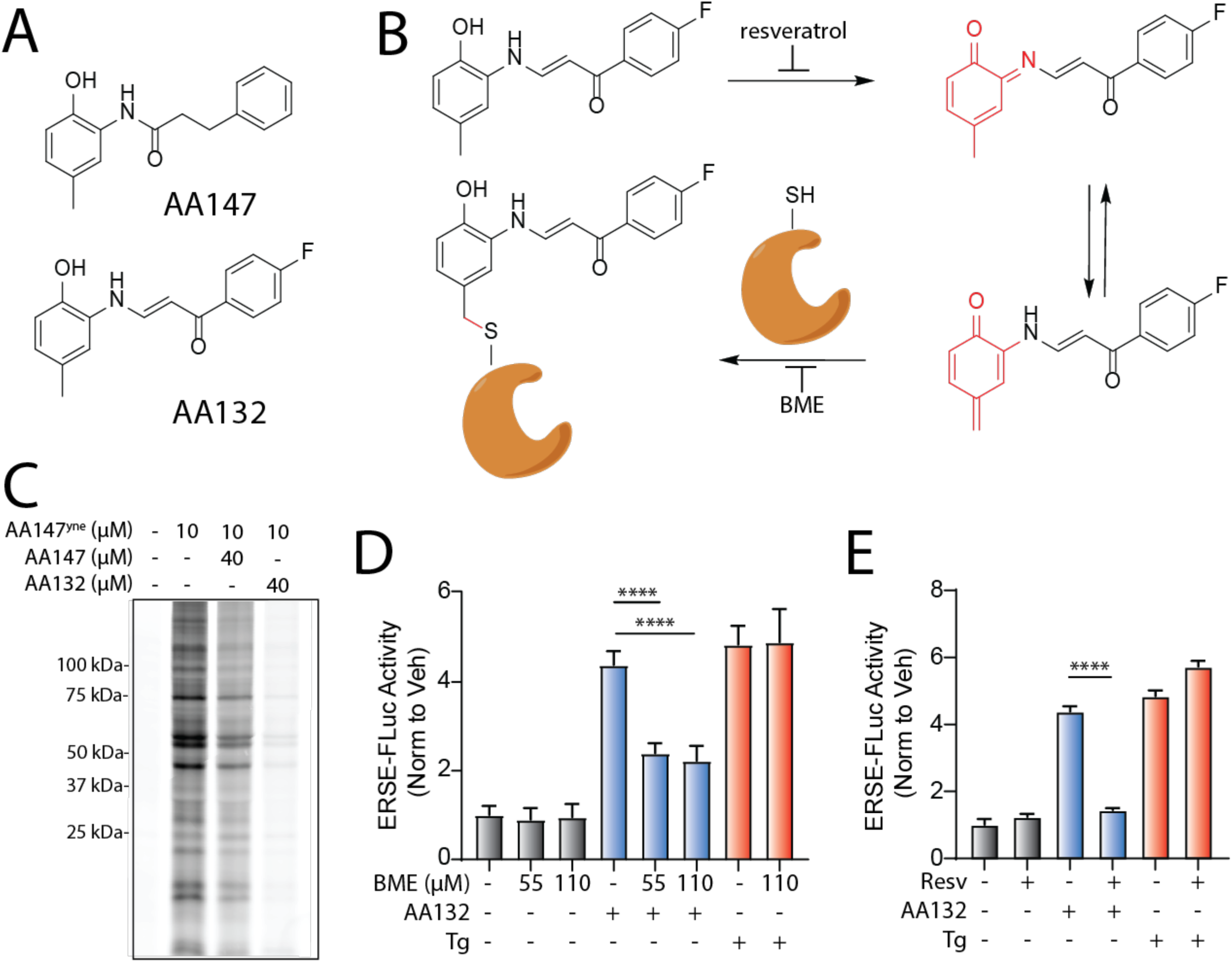
AA132 activates ATF6 signaling pathways through a mechanism involving metabolic activation and covalent protein modification. **A.** Structures of AA132 and AA147 with 2-amino-*p*-cresol substructure highlighted in red. B. Mechanism of AA132 metabolic activation and covalent protein modification C. Representative SDS-PAGE gel of Cy5-conjugated proteins from HEK293T cells treated for 4 h with vehicle (0.1% DMSO), AA147^yne^ (10 µM), the combination of AA147^yne^ (10 µM) and AA147 (40 µM), or the combination of AA147^yne^ (10 µM) and AA132 (40 µM). D. Bar graph showing the activation of the ERSE.FLuc ATF6 reporter in HEK293T cells treated with AA132 (10 µM) or thapsigargin (Tg, 500 nM) +/- β-mercaptoethanol (BME; 55 μM or 110 μM) for 18 hr. Error bars show SEM for 6 independent experiments. \*\*\**p* < 0.001, \*\*\*\**p* < 0.0001. E. Bar graph showing the activation of the ERSE.FLuc ATF6 reporter in HEK293T cells treated with AA132 (10 µM) or Tg (500 nM) +/- resveratrol (2.5 µM) for 18 hr. Error bars show SEM for 6 independent experiments. \*\*\*\**p* < 0.0001.

Intriguingly, another compound identified in our screen, (E)-1-(4-fluorophenyl)-3-((2-hydroxy-5-methylphenyl)amino)prop-2-en-1-one (AA132; **Fig 1A**), also contains the 2-amino-*p*-cresol moiety critical for AA147-dependent ATF6 activation.^27^ Despite containing the same substructure, we found that AA132 showed reduced selectivity for ATF6 transcriptional program activation in comparison to AA147; instead AA132 globally activates all three UPR signaling pathways.^19^ Understanding the requirements underlying these divergent transcriptional responses would provide insight into pharmacologic ATF6 activation and factors governing UPR activation generally. Furthermore, given the continued demonstration of the beneficial effects of selective pharmacologic ATF6 activators for ameliorating etiologically diverse diseases, such findings could guide medicinal chemistry decisions in designing more potent ATF6-selective proteostasis regulators.

In this study, we address this issue by systematically investigating the activity, selectivity, and mechanism of AA132-dependent UPR activation. We show that, like AA147, AA132 activates UPR signaling through the same mechanism observed for AA147 involving metabolic activation of the 2-amino-*p*-cresol moiety to a putative quinone methide, which then covalently modifies ER proteins, including many PDIs (**Fig 1B**). However, using a chemoproteomic probe of AA132, we demonstrate that AA132 modifies ER-resident PDIs to a greater extent than AA147. Further, we show that this observation can be attributed to the slower reactivity of the AA132-derived electrophile, allowing it to persist longer in the cellular environment prior to neutralization by endogenous nucleophiles (e.g., glutathione). This higher amount of labeling is predicted to cause ER stress, thus activating the global UPR. Consistent with this, we find that administration of AA132 at lower doses that better mimic the level of PDI modification observed with AA147 lead to selective activation of the ATF6 signaling arm of the UPR. Together, this study showcases the dynamic interplay between small chemical scaffold modifications, resulting proteome reactivity, and transcriptional selectivity for metabolically activatable ER proteostasis regulators, providing a framework for the continued development of ATF6 activating compounds for disease intervention.

## RESULTS

### AA132 activates UPR signaling through a mechanism involving metabolic activation and covalent protein modification

We previously reported that AA147 selectively activates ATF6 signaling through a process involving metabolic activation of its 2-amino-*p*-cresol substructure affording a reactive quinone methide that covalently modifies ER-localized PDIs.^27^ AA132 also contains a 2-amino-*p*-cresol moiety, suggesting that this compound could activate UPR signaling through an analogous mechanism (**Fig. 1A,B**). Consistent with this hypothesis, cotreatment of HEK293T cells with an alkyne-modified AA147 analog (AA147^yne^; 10 µM)^27^ with AA132 (40 µM) reduced AA147^yne^ protein labeling, as visualized through appending an Cy5-azide fluorophore to the terminal alkyne of conjugated proteins via Cu(I)-catalyzed azide-alkyne cycloaddition (CuAAC) (**Fig. 1C**).^32, 33^ This covalent competition reaction indicates that AA132 labels similar proteins to AA147. Intriguingly, AA132 (40 µM) showed more efficient competition with AA147^yne^ (10 µM), as compared to AA147 (40 µM), suggesting that AA132 covalently modifies target proteins more efficiently than AA147 (**Fig. 1C**).

Next, we sought to determine the sensitivity of AA132-mediated UPR activation to cotreatment with resveratrol, which blocks metabolic activation, or *β*-mercaptoethanol (BME), which reacts with the AA132-derived quinone methide prior to protein labeling (**Fig 1B**). We initially confirmed that AA132 activated luciferase-based reporters of ATF6 signaling (ERSE-FLuc)^19^, IRE1 signaling (XBP1-RLuc)^17^, and PERK signaling (ATF4-FLuc), while AA147 only significantly activated the ATF6-selective ERSE-FLuc reporter (**Fig. S1A-C**). This confirms previous results showing that AA132 activates all three signaling arms of the UPR.^19^ Cotreatment with either resveratrol or BME inhibited AA132-dependent activation of all three UPR reporters (**Fig. 1D,E**, **Fig. S1D-G**). These results support a model wherein AA132 activates global UPR signaling through a mechanism involving AA132 ER metabolic activation generating a quinone methide followed by covalent protein modification (**Fig. 1B**) – a mechanism strictly analogous to that observed for selective AA147-dependent ATF6 activation.^27^

### Synthesis and characterization of an AA132 affinity-enrichment probe to monitor protein modification

Given that AA132 cotreatment reduces AA147^yne^-dependent proteome labeling (**Fig. 1C**), we hypothesized that proteome reactivity differences may underlie the less selective transcriptional response observed with AA132 versus AA147 treatment. To test this hypothesis, we synthesized an AA132 analog with the B-ring para-fluorine replaced with an alkyne to enable subsequent affinity-enrichment experiments (**Fig. 2A**; AA132^yne^). We confirmed that AA132^yne^ activates the ATF6-selective ERSE-FLuc reporter, the IRE1-selective XBP1-RLuc reporter, and the PERK-selective ATF4-FLuc reporter (**Fig. 2B**, **Fig. S2A,B**). Further, AA132^yne^ (10 µM) induced expression of UPR target genes regulated downstream of ATF6 (*HSPA5/BiP*), IRE1 (*DNAJB9*), and PERK (*CHOP/DDIT3*) in HEK293T (**Fig. 2C**) and MEFs cells (**Fig. S2C**) at levels similar to those observed with AA132. These results further indicate that AA132^yne^, like AA132, globally activates all three arms of the UPR.

**Figure 2.**
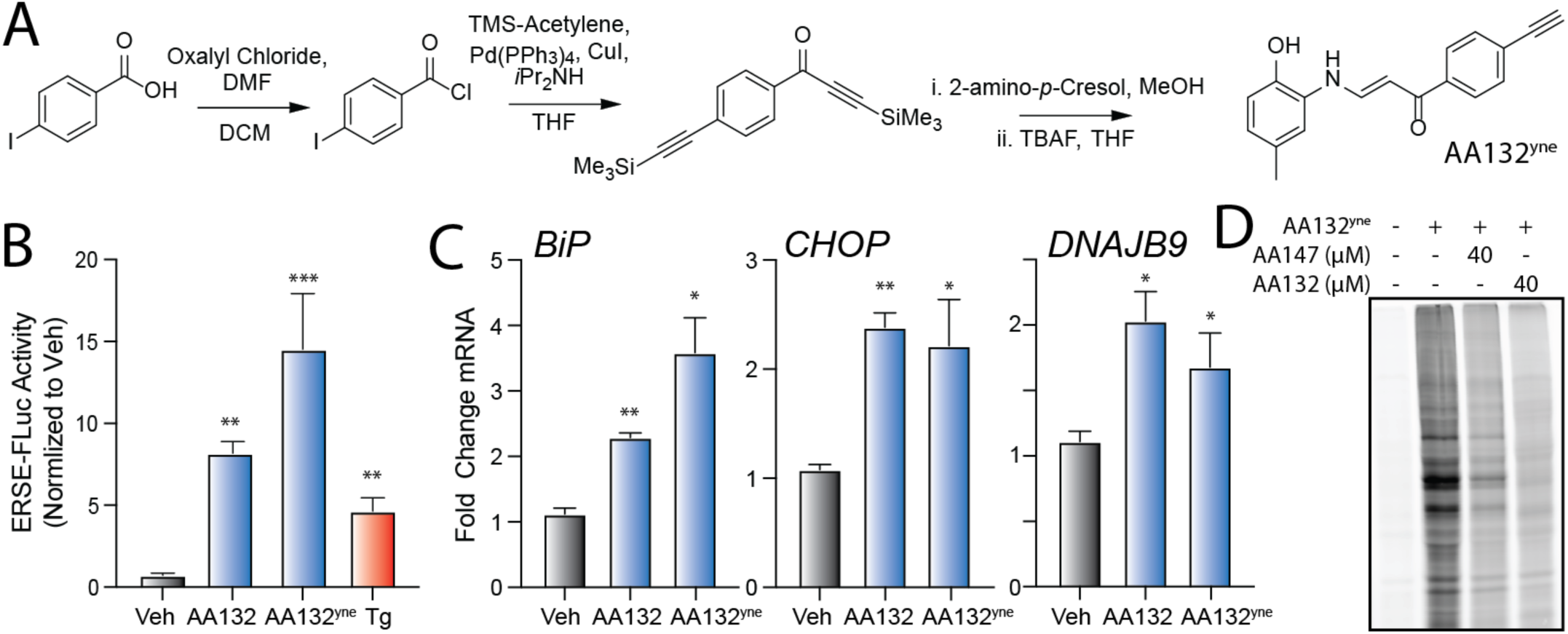
Development of a Functional Affinity Enrichment Probe for AA132. **A.** Synthetic scheme for synthesis of AA132^yne^. **B.** Bar graph showing the activation of the ERSE.FLuc ATF6 reporter in HEK293T cells treated with Veh (0.1% DMSO), thapsigargin (Tg; 500 nM), AA132(10 µM), or AA132^yne^ (10 µM) for 18 hr. \*\**p* < 0.01, \*\*\**p* < 0.001. **C.** Graph showing qPCR of the ATF6 target gene *BiP*, PERK target gene *CHOP*, and XBP1s target gene *DNAJB9* in HEK293T cells treated for 6 h with the indicated compound (10 µM). N = 3 biological replicates. \**p* < 0.05, \*\**p* < 0.01, \*\*\**p* < 0.001. **D**. Representative SDS-PAGE gel of Cy5-conjugated proteins from HEK293T cells treated for 4h with vehicle (0.1% DMSO), AA132^yne^ (10 µM), the combination of AA132^yne^ (10 µM) and AA147 (40 µM), or the combination of AA132^yne^ (10 µM) and AA132 (40 µM).

We next treated HEK293T cells with AA132^yne^ for 4h and visualized cellular protein labeling by CuAAC conjugation to a fluorescent azide–cyanine tag followed by SDS-PAGE and in-gel fluorescence scanning (**Fig 2D**).^34^ Cotreatment with four-fold excess AA147 or AA132 reduces protein labeling by AA132^yne^, indicating that these compounds target similar subsets of the cellular proteome (**Fig 2D**). As observed with AA147^yne^ (**Fig. 1C**), excess AA132 showed stronger competition with AA132^yne^ labeling than excess AA147. These results establish AA132^yne^ as an efficient probe to monitor protein conjugation afforded by oxidation of AA132 to the putative quinone methide reactive species.

#### Proteomic profiling demonstrates that AA132 preferentially modifies ER-localized proteins

Our in-gel fluorescence SDS-PAGE based competition experiments using AA147^yne^ and AA132^yne^ suggest that AA132 engages similar targets to a greater extent than AA147 (**Fig. 1C, 2D**). This suggest that AA132 possesses greater proteome reactivity, as compared to AA147. To scrutinize this hypothesis, we implemented an established affinity-purification mass spectrometry workflow to identify AA132^yne^ protein targets (**Fig. 3A**).^27^ Briefly, we treated HEK293T cells with vehicle, AA132^yne^ (10 µM), or AA132^yne^ (10 µM) with excess AA132 (40 µM) for 4 h. Diazo biotin-azide was then covalently attached to AA132^yne^-conjugated proteins by CuAAC-mediated click chemistry and the cellular protein conjugates were isolated with streptavidin affinity enrichment. Following tryptic digestion, conjugated proteins were then identified by Tandem Mass Tag (TMT)-Multi-dimensional Protein Identification Technology (MuDPIT) proteomic analysis.^35, 36^ Proteins conjugated to AA132^yne^ were further differentiated from non-specific interacting proteins using TMT reporter ion ratios between different conditions. We defined true conjugates by the following criteria: 1) a greater than 3-fold enrichment ratio from AA132^yne^-treated cells relative to DMSO treated cells, and 2) a greater than 1.5-fold reduction in enrichment ratio in cells cotreated with AA132^yne^ and excess AA132 relative to cells treated with AA132^yne^ alone (p <0.05) (**Fig. 3B** and **Table S1**). This approach identified 16 proteins modified by AA132^yne^, 10 of which were previously identified targets of AA147^yne^ (**Fig. 3C**).^27^ Intriguingly, like AA147^yne^, AA132^yne^ preferentially conjugated to ER-localized proteins (11/16, 69% of identified targets are localized to the ER; **Fig. S3A**).^27^ Further, the most abundant members of the PDI family, including PDIA3, PDIA4, PDIA6, PDIA1, and TXNDC5, are well-represented among the shared targets of AA132^yne^ and AA147^yne^. These results indicate that AA132^yne^, like AA147^yne^, selectively modifies ER-resident proteins, most notably PDIs, likely reflecting the similar physicochemical properties between AA147 and AA132 (**Fig. S3B**). This also further supports our model whereby AA132^yne^ activates UPR signaling through a mechanism involving covalent modification of ER-localized PDIs (**Fig. 1B**).

**Figure 3.**
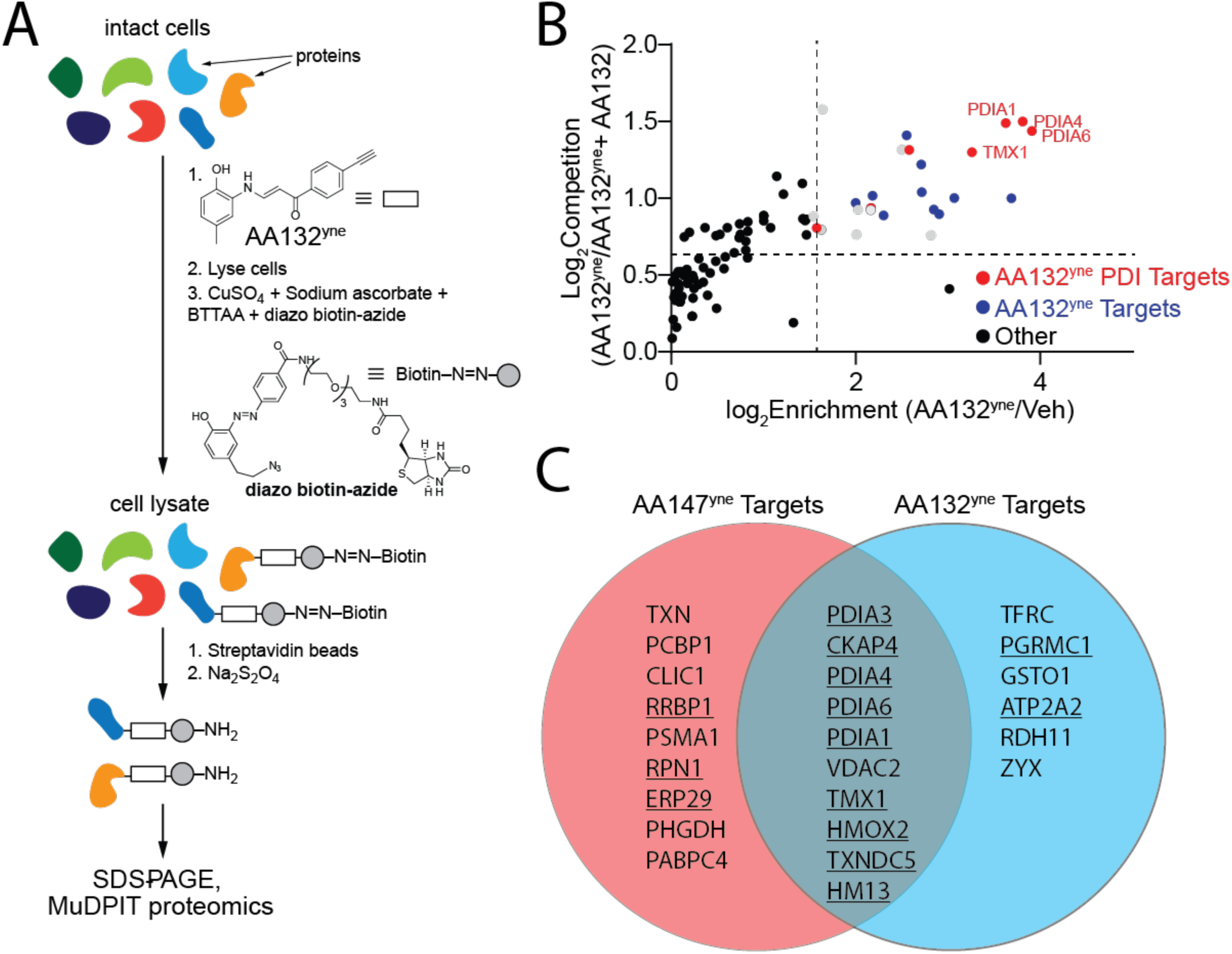
AA132^yne^ Covalently Modifies ER PDIs. **A.** Schematic showing the protocol for affinity purification of proteins covalently modified by AA132^yne^. Adapted from Paxman et al., 2018. **B.** Plot showing Log2 Fold Change of AA132^yne^-enriched proteins relative to vehicle (X-axis) and Log2 Fold Change of competition ratio (AA132^yne^ + Veh / AA132^yne^+ AA132) (Y-axis). Dotted lines indicate significantly enriched proteins (>3-fold; x-axis) and proteins with a significant competition ratio (>1.5 fold; y-axis). Red circles identify AA132^yne^ targets with PDI GO annotation; blue circles identify additional AA132^yne^ targets identified across two separate experiments. Grey circles identify AA132^yne^ targets identified in one experiment. Data included in **Table S1**. **C.** Venn Diagram showing unique targets of AA147^yne^ (red) or AA132^yne^ (blue). Underlined proteins have the GO annotation GO:0044432 “*Endoplasmic Reticulum Part”*.

### AA132 shows enhanced PDI labeling as compared to AA147

Although AA132 and AA147 label similar ER PDI family members, these two compounds induce distinct transcriptional profiles, with AA147 showing selective ATF6 activation and AA132 showing activation of all three arms of the UPR.^19^ To better define this discrepancy, we performed quantitative TMT-MuDPIT proteomics to directly compare the populations of proteins labeled by AA147^yne^ versus AA132^yne^ in HEK293T cells (**Table S2**). Despite labeling similar proteins (**Fig. 4A**), we found that AA132^yne^ showed higher overall labeling of multiple targets, as compared to AA147^yne^ at the same concentration (10 µM) (**Fig. 4B**). This includes greater labeling for numerous PDIs linked to ATF6 activation, including PDIA1, PDIA3, PDIA4, PDIA6, and TXNDC5 (**Fig. 4A,B**). Similar results were observed in other cells types, including liver-derived HepG2 cells (**Fig. S4A,B**, **Table S2**). We confirmed the increased labeling of PDIA3 and PDIA4 in HEK293T cells by biotin conjugation, streptavidin enrichment, and quantitative immunoblotting (**Fig. S4C**). These results suggest that increased modification of PDIs could define the differential transcriptional signaling observed between AA147 and AA132.

**Figure 4.**
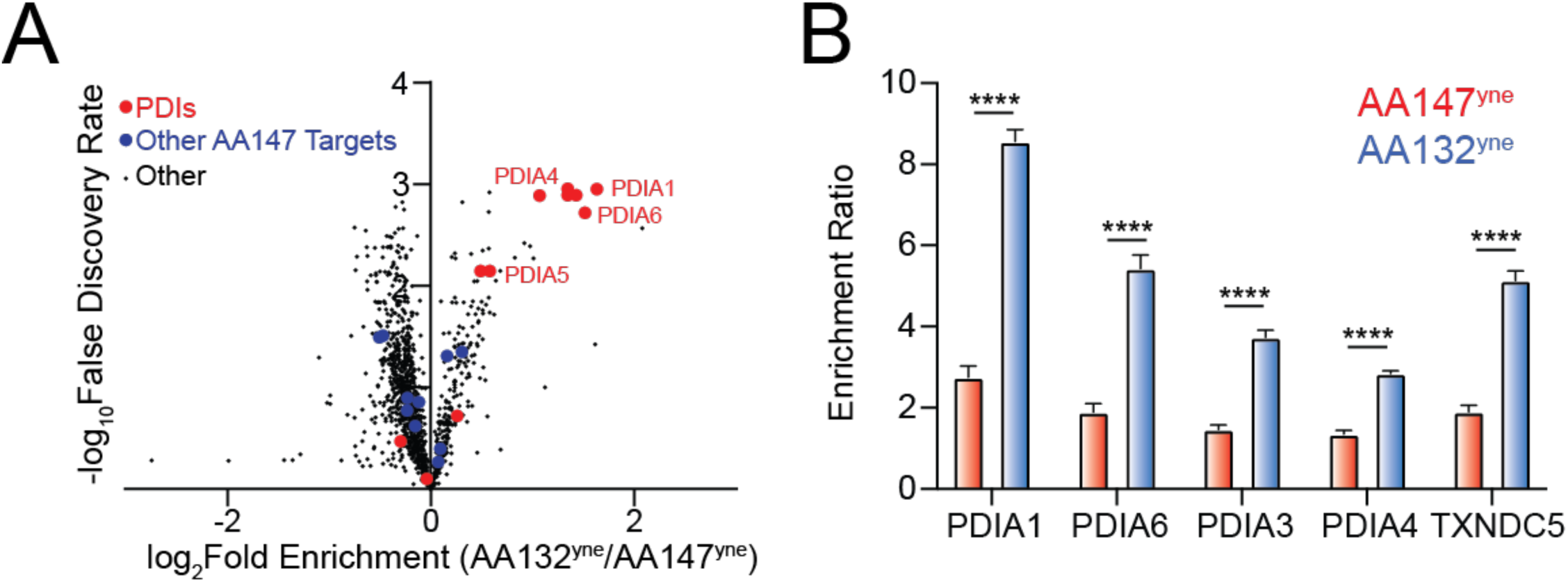
AA132^yne^ Shows Higher PDI Labeling as Compared to AA147^yne^. **A.** Volcano plot showing log2 fold enrichment of AA132^yne^ labeled proteins relative to AA147^yne^ labeled proteins (x-axis) versus the –log FDR (y-axis) in HEK293T cells (10 µM, 6h). Proteins with GO annotation for PDI (GO:0003756) labeled in red and additional previously defined AA147^yne^ targets labeled in blue. Data shown in **Table S2 B.** Bar Graph of TMT reporter ion enrichment ratio of select PDIs from data shown in Fig. 4A in HEK293T cells treated with the indicated compound relative to DMSO (N = 4 biological replicates). ****p < 0.001 for a multiple unpaired t-test.

### AA132^yne^ shows slower labeling kinetics as compared to AA147^yne^

Given the relative structural similarity between AA147 and AA132, we sought to identify the basis for the divergence in proteome labeling and subsequent transcriptional selectivity. We initially predicted that the small structural differences between AA147 and AA132 could lead to changes in the stability of target proteins upon small molecule conjugation. However, we did not see significant differences in resistance to heat denaturation for PDIA1 labeled with AA147 or AA132 (**Fig. S5A,B**).

We next tested whether differences in the kinetics of conjugate formation by AA147 or AA132 could mediate their differential proteome labeling. We performed an in-gel fluorescence time course experiment in HEK293T cells monitoring AA147^yne^ or AA132^yne^ conjugate formation. Intriguingly, despite AA132^yne^ showing higher overall PDI labeling after 6h of treatment (**Fig. 4A,B**), AA132^yne^ showed slower kinetics of proteome labeling, as compared to AA147^yne^ (**Fig. 5A-C**). The relative extent of PDI labeling by AA132 and AA147 becomes equal after four hours, suggesting that AA132 has a longer operational half live *in cellulo* (**Fig. 5C**). Similar results were observed in other cell types, including ALMC2 plasma cells (**Fig. S5C,D**). This suggests that AA132^yne^ labels proteins at a slower rate than AA147^yne^. If correct, this effect is predicted to make AA132^yne^ protein labeling more susceptible to inhibition afforded by exogenous nucleophiles such as BME. Consistent with this, cotreatment with BME showed greater inhibition of AA132^yne^ labeling, as compared to AA147^yne^ labeling (**Fig. S5E,F**). Together, these results indicate that AA132^yne^ forms protein conjugates slower, as compared to AA147^yne^, despite ultimately reaching higher levels. This suggests that AA132^yne^ forms a more stable, longer-lived electrophilic species in its mechanism of protein conjugation.

**Figure 5.**
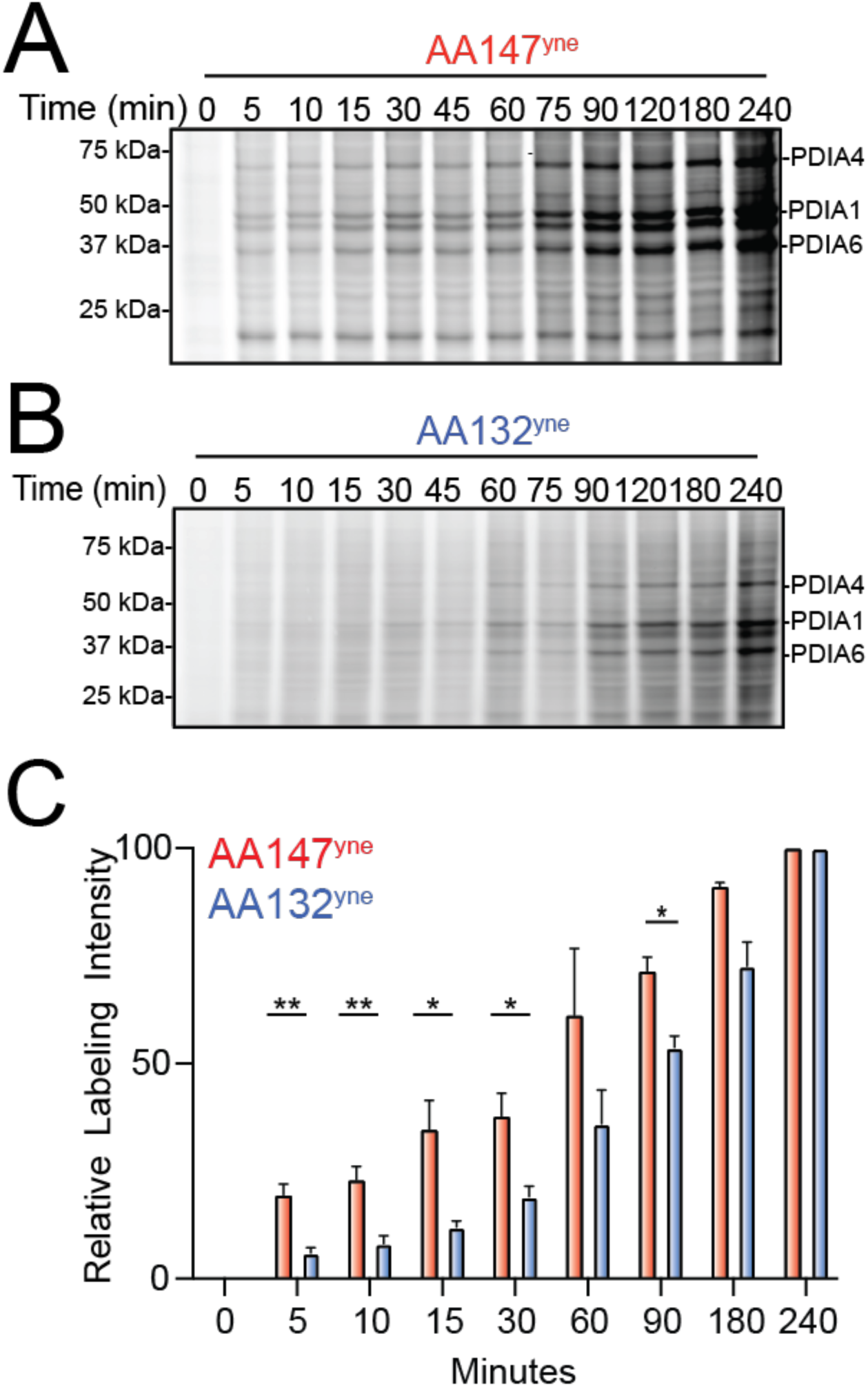
AA132^yne^ Shows Slower Protein Labeling Kinetics as Compared to AA147^yne^. **A.** Representative SDS-PAGE gel of Cy5-conjugated proteins from HEK293T cells treated for the indicated time point with AA147^yne^ (10 µM). **B.** Representative SDS-PAGE gel of Cy5-conjugated proteins from HEK293T cells treated for the indicated time point with AA132^yne^ (10 µM). **C.** Quantification of gels described in Fig 5A and Fig 5B. Y-axis labeling relative intensity represents PDIA4 intensity normalized to labeling at 6h. Error bars represent S.E.M (N = 4 Biological Replicates). \**p* < 0.05, \*\**p* < 0.01.

#### Dose-dependent regulation of AA132 transcriptional selectivity

AA132 and AA147 induce distinct transcriptional profiles in HEK293T cells after 6h treatment. AA147 selectively activates the ATF6 arm of the UPR, while AA132 activates all three arms of the UPR.^19^ Our results indicate that the distinct transcriptional profiles induced by these two compounds could be attributed to differences in PDI labeling, with AA132 modifying PDIs to a greater extent than AA147 (**Fig. 4B**). This would suggest that decreasing AA132-dependent PDI modification should result in increased transcriptional selectivity for ATF6 activation. To test this, we monitored mRNA expression by RNAseq in HEK293T cells treated with increasing concentrations of AA132 from 0.1 - 30 µM for 6 h (**Table S3**). We confirmed dose-dependent protein modification in HEK293T cells treated with increasing concentrations of AA132 (**Fig. S6A**). As expected, the number of differentially expressed genes (DEGs) increased with increasing AA132 concentrations (**Fig. 6A**). Gene ontology (GO) analysis of significantly induced genes in AA132-treated cells showed selective increases in terms associated with ER function, ER stress, and the UPR (**Fig. 6B,C**, **Fig. S6B-D**, **Table S4**). These results confirm the genome wide transcriptional specificity of AA132 for UPR activation reported previously.^19^

**Figure 6.**
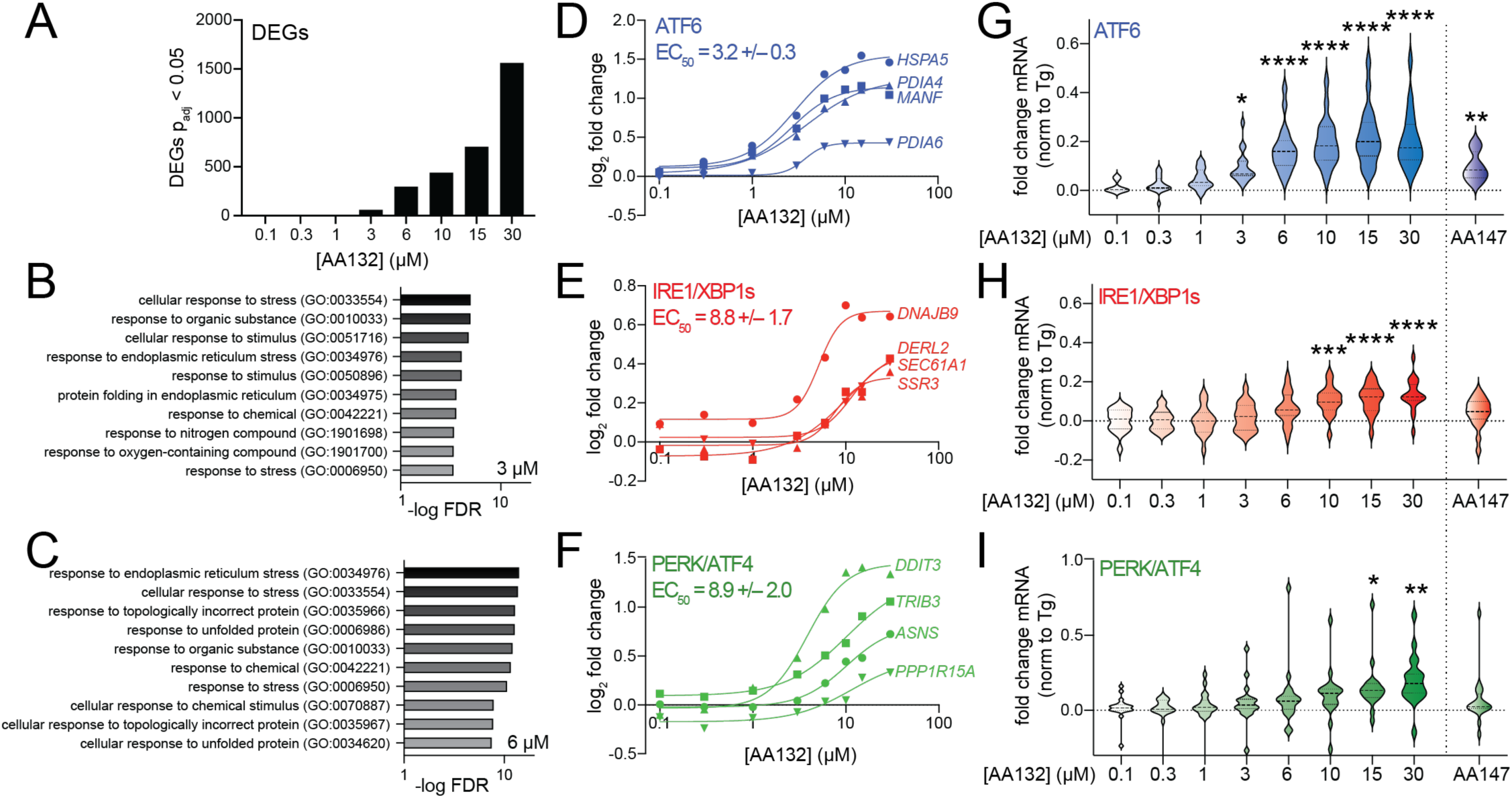
AA132 Selectively Activates ATF6 Transcriptional Signaling at Lower Doses. **A.** Differentially expressed genes (padj<0.05) from RNAseq data of HEK293T cells treated with vehicle or the indicated dose of AA132 for 6 h (N = 3 replicates per condition). RNAseq data is included in **Table S3. B,C**. Top-10 GO terms for significantly induced genes (fold change>1.3, p<0.05) identified by RNAseq in HEK293T cells treated with 3 µM (**B**) or 6 µM (**C**) AA132 for 6 h. RNAseq data is included in **Table S3**. Full GO analysis is included in **Table S4**. **D-F**. Fold change, relative to vehicle, for select ATF6 target genes (**D**), IRE1/XBP1s target genes (**E**), or PERK/ATF4 target genes (**F**) from RNAseq data of HEK293T cells treated with increasing concentrations of AA132 for 6 h. The average EC50 for the four target genes representing each UPR pathway are shown. **G-I**. Fold change, relative to vehicle, for genesets of 15-20 genes regulated downstream of ATF6 (**G**), IRE1/XBP1s (**H**), or PERK/ATF4 (**I**) from RNAseq of HEK293T cells treated with increasing concentrations of AA132 for 6 h. The fold change expression of individuals genes was normalized to that observed with the global ER stressor thapsigargin (Tg), as reported previously.^19, 37^ The impact of AA147 (10 µM; 6 h) on these genesets is shown on the right. The expression of UPR genesets is shown in **Table S5**. *p < 0.05, **p < 0.01, ***p < 0.005, ****p < 0.001 for one-way ANOVA.

Next, we sought to define the transcriptional selectivity of increasing concentrations of AA132 for activation of the three arms of the UPR. Monitoring expression of select target genes of ATF6 (*HSPA5*, *PDIA4*, *PDIA6, MANF*), IRE1/XBP1s (*DNAJB9, SSR3, SEC61A, DERL2*), and PERK/ATF4 (*DDIT3, PPP1R15A, ASNS, TRIB3*) shows dose-dependent increases of these genes (**Fig. 6D-F**). Interestingly, the EC50 for ATF6-target genes (3.2 µM) is less than that observed for IRE1/XBP1s and PERK/ATF4 target genes (8.8 and 8.9 µM, respectively). This suggests that ATF6 signaling is induced at lower concentrations of AA132, as compared to IRE1/XBP1s and PERK/ATF4. Further, normalizing the expression of 15-20 target genes regulated by ATF6, IRE1/XBP1s, or PERK/ATF4 observed following 6h treatment with AA132 to that observed with the global ER stressor thapsigargin (Tg) shows that ATF6 target genes are globally and significantly induced starting at 3 µM, while IRE1/XBP1s and PERK/ATF4 target genesets are significantly induced starting at 10 and 15 µM, respectively (**Fig. 6G-I**, **Table S5**).^37^ Interestingly, the activation of these genesets observed at 3 µM AA132 match the selective ATF6 activation observed in HEK293T cells treated with AA147 (10 µM, 6 h; **Fig. 6G-I**, right). These results support a model whereby lower concentrations of AA132 (3-6 µM) selectively activate ATF6 transcriptional signaling, while higher concentrations (>10 µM) lead to global UPR activation.

## DISCUSSION

Here, we sought to define the molecular basis for the differential transcriptional selectivity of AA147 and AA132 with regards to arm-selective UPR activation. Like AA147, we show that AA132 achieves pharmacologic UPR modulation through a mechanism involving metabolic oxidation of its 2-amino-*p*-cresol moiety yielding a putative quinone methide that subsequently covalently modifies ER-localized proteins, most notably multiple PDIs.^27^ Despite sharing numerous protein targets, comparative chemoproteomic experiments revealed that AA132 labels ER proteins, including PDIs, to a greater extent than AA147 at treatment times exceeding four hours. This divergence in proteomic reactivity provides a mechanism to explain the differential selectivity for UPR activation observed upon AA132 or AA147 treatment, with AA132 activating all three arms of the UPR and AA147 selectively activating ATF6 UPR signaling.^19^ Our results indicate that this greater PDI labeling is attributed to increased stability of the metabolically activated AA132 compound, increasing its diffusion within the cell and subsequent accessibility to a larger pool of ER proteins. The increased labeling of ER proteins afforded by AA132 provides a mechanism to explain the global activation of UPR signaling pathways observed with this compound, as higher labeling would be predicted to further disrupt PDI activity and subsequently increase UPR signaling, possibly by producing misfolded secretory proteins. In contrast, the more modest PDI inhibition induced by AA147 would only be sufficient to promote ATF6 reduction, monomerization, and trafficking, leading to selective activation of ATF6 transcriptional activity. Consistent with this, lower concentrations of AA132 decrease PDI modifications and lead to selective ATF6 activation, mimicking the transcriptional selectivity observed with AA147. Together, these results highlight the connection between electrophilic reactivity, apparent quinone methide half-life, and transcriptional selectivity for pharmacologic ATF6 activators, establishing new opportunities to develop next generation compounds of this class with improved activity and selectivity.

While we and others have demonstrated the importance of PDI family members in dictating ATF6 activity, the PDIs also have critical roles in regulating the IRE1 and PERK arms of the UPR. PDIA6 interacts with luminal domains of both PERK and IRE1 to prevent hyperactivation of these UPR pathways.^38^ Similarly, both PDIA1 and PDIA3 are implicated in regulating PERK signaling in cancer cells in the presence or absence of ER stress.^39^ In addition, other PDIs, including PDIA5 and ERP18, interact with ATF6 luminal domain disulfides to regulate ATF6 anterograde trafficking to the Golgi prior to proteolytic activation.^28–31^ Previous results indicate that AA147 activates ATF6 signaling through selective modification of only a subset of specific ER PDIs. For example, AA147 was shown to modify only ∼20% of PDIA1 within the ER.^27^ This suggests that the ability for AA147 to selectively activate ATF6 signaling lies in its unique capacity to modify a small population of specific PDIs. Consistent with this, genetic depletion of PDIs including *PDIA1*, *PDIA3*, *PDIA4*, and *PDIA5* limit AA147-dependent ATF6 activation.^27^

Our results suggest that the increased modification of PDIs afforded by AA132 underlies its ability to globally activate UPR signaling. We show that AA132 modifies multiple PDIs to a greater extent than AA147, including PDIA1 and PDIA6. Interestingly, highly selective covalent PDIA1 inhibitors that modify PDIA1 to greater extents than AA147 can activate IRE1/XBP1s signaling.^40, 41^ This suggests that increased targeting of PDIA1 by AA132 may contribute to the IRE1/XBP1s activation observed under these conditions. Similarly, increased PDIA6 modification could mitigate the repression of IRE1 and PERK hyperactivity afforded by this PDI ^38^, increasing activity of these two UPR signaling pathways. To support this model, we show that decreasing AA132-dependent modifications of PDIs, including of PDIA1 and PDIA6, decreases activation of IRE1 and PERK transcriptional signaling and allows selective ATF6 transcriptional activity. This suggests that the transcriptional activity of AA132 for global UPR activity can be viewed as a graded disruption in the tightly controlled regulatory roles of ER PDIs involved in regulating ER proteostasis and UPR signaling. While we cannot rule out different protein targets contributing to the activation of different arms of the UPR, our data are consistent with a more pronounced on-target mechanism involving increased PDI modification being responsible for the global UPR activating capacity of AA132.

AA132 and AA147 have close structural similarity, with both compounds containing the 2-amino-*p*-cresol substructure critical for compound activity. However, there exists a profound difference in both transcriptional activity and equilibrium proteome labeling between AA147 and AA132. A possible modulatory role of the linker region on the activity of the metabolically derived electrophile helps reconcile these opposing results. AA132 contains an enaminone linkage, while AA147 contains an amide linkage. Increased electrophile stability or decreased rate of metabolic activation mediated by the amide to enaminone substitution could explain the observed divergence in biological effects. For instance, activation of AA132 to the requisite electrophilic species may require additional isomerization steps, possibly involving the enaminone moiety, in an alternate metabolic activation pathway.^42, 43^ The rate of tautomerization of quinone imine/quinone methide moieties, by nonenzymatic or enzymatic means, may be modulated by the linker region.^44^ In addition, π-conjugation present in the enaminone class of metabolically activatable proteostasis regulators may stabilize the resulting electrophilic species, and thus enable a larger sphere of activity.^45^ Importantly, the PDIs are not amongst the most reactive ER-resident cysteines, and thus increased labeling is not likely to be a simple function of electrophile reactivity towards hyperactive cysteines.^46^ Nevertheless, other soft electrophilic species similar to those derived from AA147 or AA132, such as quinone-based electrophiles, have preferential reactivity towards the PDIs.^47–50^ Our kinetic analyses observed in context with the greater reduction in AA132^yne^ conjugation seen with BME cotreatment suggest that increased stability, rather than increased reactivity, may underlie our observation of greater PDI engagement with the enaminone class of ER proteostasis regulators. Finally, the identification of unique ER AA132^yne^ targets not seen with AA147^yne^ could be a result of a larger diffusion radius from site of AA132^yne^ electrophilic metabolite generation. Alternatively, subtle reactivity differences between AA132^yne^- and AA147^yne^-derived electrophiles may differentially facilitate conjugation with previously annotated reactive cysteines on some of these targets.^51–53^

Our chemoproteomic results provide insight into the narrow window of PDI conjugation necessary for selective ATF6 activation mediated by metabolically activatable ER proteostasis regulators. In this view, the activity of a few of the PDIs can be viewed as a rheostat controlling a spectrum of UPR transcriptional activities. From a therapeutic perspective, concomitant activation of ATF6 and IRE1 signaling leads to a uniquely remodeled ER proteostasis environment preferable for some disease contexts, as compared to individual activation of individual UPR pathways.^54, 55^ Thus, the extent of global UPR activation observed with AA132 may be of some value in certain disease contexts. Regardless, a continued understanding of the factors driving selectivity of the transcriptional response induced by ER proteostasis regulators is essential for the development of improved ATF6 activators for treatment of etiologically diverse diseases.

## ACKNOWLEDGEMENTS

We thank Emily P. Bentley (Scripps Research) for critical reading of this manuscript. Funding for this work was provided by the National Institutes of Health (AG046495 to RLW, JWK) and the Skaggs Institute for Chemical Biology (JWK).

## SUPPLEMENTARY TABLE LEGENDS

**Table S1.** Excel spreadsheet showing the enrichment and competition ratio for proteins identified as targets of AA132^yne^. Related to **Fig. 3**.

**Table S2.** Excel spreadsheet showing fold enrichment for AA132^yne^/AA147^yne^ of proteins identified in proteomics experiments performed in HEK293T or HepG2 cells. Related to **Fig. 4** and **Fig. S4**.

**Table S3.** Excel spreadsheet showing DESeq outputs for HEK293T cells treated with increasing doses of AA132 or AA147. Related to **Fig. 6**.

**Table S4.** Excel spreadsheet showing GO analysis for RNAseq data for HEK293T cells treated with the indicated concentration of AA132. Related to **Fig. 6**.

**Table S5.** Excel spreadsheet showing the expression of transcriptional targets of ATF6, IRE1/XBP1s and PERK signaling from RNAseq data of HEK293T cells treated with the indicated concentration of AA132 or AA147. Related to **Fig. 6**.

## MATERIALS AND METHODS

### Cell Culture

HEK293T-Rex (ATCC), HEK293T (ATCC), and HepG2 (ATCC) were cultured in high-glucose Dulbecco’s Modified Eagle’s Medium (DMEM) supplemented with glutamine, penicillin/streptomycin and 10% fetal bovine serum (FBS). Cells were routinely tested for mycoplasma every 6 months. We did not further authenticate the cell lines. All cells were cultured under typical tissue culture conditions (37 °C, 5% CO2). ALMC-2 (a kind gift from Diane Jelinak’s laboratory) were cultured in Iscove’s Modified Dulbecco’s Medium (IMDM) GlutaMAX (Life Technologies) supplemented with penicillin/streptomycin, 5% fetal bovine serum and 2 ng/mL interleukin-6 (IL-6). All cells were cultured under typical tissue culture conditions (37°C, 5% CO2)

### Measurement of UPR activity using luciferase reporters

HEK293T-Rex cells expressing the ERSE.FLuc^19^, XBP1s.RLuc ^17^, or ATF4.FLuc reporter were plated at 100 µL/well from suspensions of 200,000 cells/mL in white clear-bottom 96-well plates (Corning) and incubated at 37°C overnight. The following day, cells were treated with 25 µL of compound-containing media to give final concentration as described before incubating for 18 hr at 37°C. The plates were equilibrated to room temperature, then either 125 μL of Firefly luciferase assay reagent-1 (ERSE.FLuc and ATF4.FLuc) or Renilla luciferase assay reagent-1 (XBP1s.RLuc) (Targeting Systems) were added to each well. Samples were dark adapted for 10 min to stabilize signals. Luminescence was then measured in an Infinite F200 PRO plate reader (Tecan) and corrected for background signal (integration time 250 ms). All measurements were performed in biologic triplicate.

### Quantitative RT-PCR

The relative mRNA expression levels of target genes were measured using quantitative RT-PCR. Cells were treated as described at 37°C, harvested by trypsinization, washed with Dulbecco’s phosphate-buffered saline (GIBCO), and then RNA was extracted using the QuickRNA Miniprep Kit (Zymo). qPCR reactions were performed on cDNA prepared from 500 ng of total cellular RNA using the High-Capacity cDNA Reverse Transcription Kit (Applied Biosystems). PowerSYBR Green PCR Master Mix (Applied Biosystems), cDNA, and appropriate primers purchased from Integrated DNA Technologies (see Table below) were used for amplifications (6 min at 95°C, then 45 cycles of 10 s at 95°C, 30 s at 60°C) in an ABI 7900HT Fast Real Time PCR machine. Primer integrity was assessed by a thermal melt to confirm homogeneity and the absence of primer dimers. Transcripts were normalized to the housekeeping genes RPLP2 and all measurements were performed in biological triplicate. Data were analyzed using the RQ Manager and DataAssist 2.0 software (ABI). qPCR data are reported as mean ± standard deviation plotted using Prism GraphPad.

#### Sequences of Primers for qPCR.

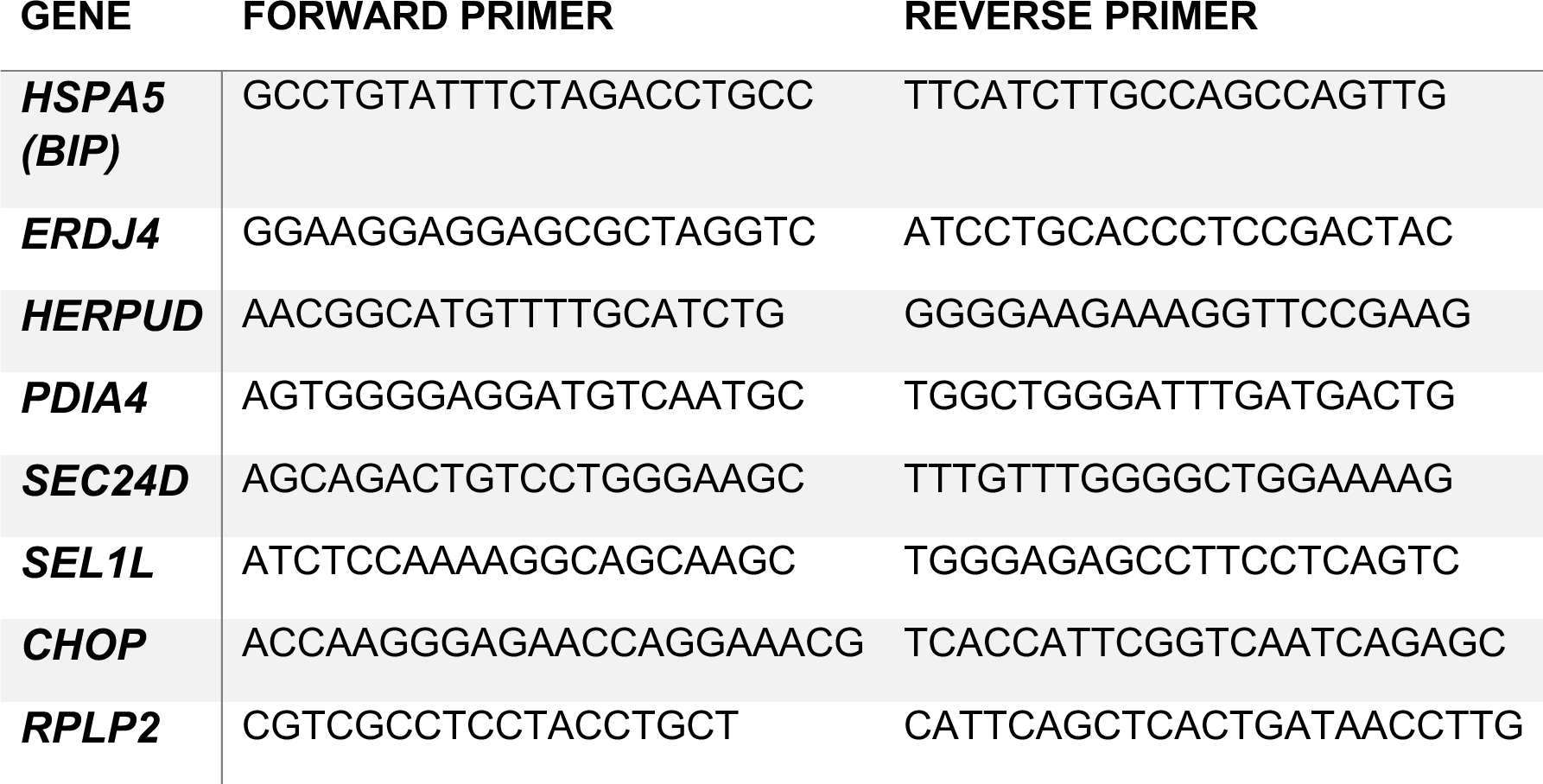

### SDS-PAGE In-Gel Fluorescence Scanning

ALMC-2 cells (200,000 cells/mL) or HEK293T cells (250,000 cells/well) were treated with indicated compound in six-well plates at indicated concentration and time period. Cells were lysed in radioimmunoprecipitation assay (RIPA) buffer (150 mM NaCl, 50 mM Tris pH 7.5, 1% Triton X-100, 0.5% sodium deoxycholate, and 0.1% SDS) supplemented with fresh protease inhibitor cocktail (Roche, Indianapolis, IN) and centrifuged for 20 min at 16000×g following a 30-minute incubation. Protein concentration of supernatant determined by BCA assay (Thermo Fisher) and normalized to give 42.5 µL at 2.35 mg/mL (100 µg/total protein). 7.5 µL ‘click chemistry master mix’ was added to each sample to give final concentrations of 100 µM of Cy5-azide (Click Chemistry Tools, Scottsdale, AZ), 800 µM copper (II) sulfate, 1.6 mM BTTAA ligand (2-(4-((bis((1-tert-butyl-1H-1,2,3-triazol-4-yl)methyl)amino)methyl)-1H-1,2,3-triazol-1-yl)acetic acid) (Albert Einstein College), and 5 mM sodium ascorbate. Reaction incubated at 30° C for 1 hour while shaking before CHCl3/MeOH protein precipitation. Dried protein was redissolved in 20 µL 1X SDS loading buffer and 25 µg was loaded on gel for SDS-PAGE in-gel fluorescence scanning and subsequent visualization using an Odyssey Infrared Imaging System (Li-Cor Biosciences).

### Immunoblotting of AA132^yne^ or AA147^yne^ conjugated proteins

HEK293T cells grown to 80-90% confluency in 10 cm plates were treated with 10 µM indicated compound (AA147^yne^, AA132^yne^, or Veh) for 6h at 37 °C. The cells were washed with PBS before harvesting with tryspin, pelleting (500 g, 5 min), and washed with PBS (1 mL). Cells pellets were resuspended in radioimmunoprecipitation assay (RIPA) buffer before sonication with a probe tip sonicator to lyse the cells (15 sec, 3 sec on/2 off, 30% amplitude). The lysates were cleared via centrifugation and the concentration of protein adjusted to 4 mg / mL using the BCA assay. 2 g protein (500 µL) was taken and reacted with a mixture of diazo biotin azide (100 µM), copper (II) sulfate (800 µM), BTTAA (1.6 mMM), sodium ascorbate (5.0 mM) for 90 min at 30 °C with shaking (600 rpm). The reaction was quenched with the sequential addition of cold methanol (4x volume), chloroform (1x volume), and DPBS (4x volume) to precipitate proteins. Proteins were pelleted by centrifugation (4,700 g, 10 min, 4 °C). The supernatant was discarded, and the pellets dried under air for 5 min. Protein pellets were resuspended in 6M urea in PBS (500 µL) with brief sonication. 25 µL of sample was taken as the “input” for Western blot analysis. Sample diluted with 5.5 mL DPBS (.2% SDS) and streptavidin agarose resin (100 µL, washed 3 x 1 mL with PBS) was added to each sample before incubation for 18 h at 24 °C with rotation. The beads were pelleted via centrifugation (3000 g, 2 min) and washed with 0.2% SDS in DPBS, DPBS (2x 2 mL). 100 µL freshly prepared 50 mM sodium dithionite added to beads to cleave protein conjugates and incubated at 30 ° C for 1h. Supernatant transferred to new 2 mL Eppendorf and precipitated with cold methanol (4x volume), chloroform (1x volume), and DPBS (4x volume). Proteins resuspended in 1x SDS-PAGE gel loading buffer (20 µL) and heated for 5 min at 95°C prior to separation via SDS-PAGE and transfer to PVDF membranes for immunoblot analysis of indicated proteins (rabbit PDIA4 (1:2000) (Protein Tech), rabbit PDIA3 (1:1000) (Protein Tech), mouse GAPDH (1:1000) (Cell Signaling), Goat anti-rabbit IRdye 800-cw (Licor) (1:10000), Goat anti-mouse IRdye-680RD (Licor) (1:10000).

### Cellular Thermal Shift Assay

ALMC2 cells grown to concentration of 2 million cells/mL and 15 mL cell suspension incubated in T75 flask with indicated compound (10 µM, 2h, 37° C). After treatment, cells were pelleted by centrifugation (3 min, 300 g) and washed with PBS before resuspension in PBS at 30 million cells/mL. 100 µL cell suspension was added to 0.2 mL PCR tube before heat treatment at indicated temperature for 3 min and room temperature incubation for 3 min. Samples snap frozen at -80 C were lysed by sequential freeze-thaw cycles and centrifuged (15,000 g, 10 min) to pellet insoluble material. Soluble fractions were boiled for 5 min in Laemmli buffer with 100 mM DTT before loading onto SDS-PAGE gels. Proteins were transferred from gel slabs to PVDF membranes and blotted using rabbit anti-PDIA1 antibody (1:1000) Protein Tech) and visualized on the Odyssey Infrared Imaging System (Li-Cor Biosciences).

### Chemoproteomic Analysis

HEK293T cells in 10 cm plates at 80-90% confluency were treated for 6 h with vehicle (0.1% DMSO), AA132^yne^ (10 μM), or the combination of AA132^yne^ (10 μM) and AA132 (40 μM) at 37 °C. The cells were washed with PBS before harvesting with tryspin, pelleting (500 g, 5 min), and washed with PBS (1 mL). Cells pellets were resuspended in radioimmunoprecipitation assay (RIPA) buffer before sonication with a probe tip sonicator to lyse the cells (15 sec, 3 sec on/2 off, 30% amplitude). For each sample, 1 g lysate (500 µL) were reacted with click reagents to give final concentrations as follows: 100 µM of diazo biotin-azide (Click Chemistry Tools, Scottsdale, AZ), 800 µM copper (II) sulfate, 1.6 mM BTTAA ligand (2-(4-((bis((1-tert-butyl-1H-1,2,3-triazol-4-yl)methyl)amino)methyl)-1H-1,2,3-triazol-1-yl)acetic acid) (Albert Einstein College), and 5 mM sodium ascorbate. The reaction was placed on a shaker at 1000 rpm at 30 °C for 90 The reaction was quenched with the sequential addition of cold methanol (4x volume), chloroform (1x volume), and DPBS (4x volume) to precipitate proteins. Proteins were pelleted by centrifugation (4,700 g, 10 min, 4 °C). The supernatant was discarded, and the pellets dried under air for 5 min. Protein pellets were resuspended in 6M urea in PBS (500 µL) with brief sonication. 50 µL of high-capacity streptavidin beads were washed with PBS and mixed with the protein solution in 6 mL of phosphate-buffered saline (PBS). This suspension was placed on a rotator or a shaker and agitated for 2 h. The beads were centrifuged and washed 5 times with PBS and 1% SDS. The protein was eluted from the beads by two washes of 50 mM sodium dithionite in 1% SDS for 1 h and then precipitated by chloroform/methanol precipitation as described above. 50 µL of freshly made 1:1 mixture 200 mM TCEP·HCl in DPBS and 600 mM K2CO3 in DPBS was added to each sample before incubation at 37°C for 30 minutes while shaking. Alkylation of reduced thiols was achieved by addition of 70 µL freshly prepared 400 mM iodoacetamide in DPBS and incubation at room temperature while protected from light. The reaction was quenched by adding 130 µL of 10% SDS in DPBS and then diluted to approximately 0.2% SDS via DPBS (5.5 mL) and incubated with preequilibrated streptavidin agarose beads (3x1 mL PBS wash). The samples were rotated at room temperature for 1.5 hours, centrifuged at 2000 rpm for 2 minutes, and then washed sequentially with 5 mL 0.2% SDS in DPBS, 5 mL DPBS, and 5 mL 100 mM TEAB (Thermo Cat #90114) pH 8.5 to remove non-binding proteins. The beads were transferred to low-bind 1.5 mL Eppendorf tubes and the bound proteins digested overnight at 37 °C in 200 µL 100 mM TEAB containing 2 µg sequencing grade porcine trypsin, 1 mM CaCl2, and 0.01% ProteaseMax (Promega Cat #V2071). The beads were centrifuged at 2000 rpm for 5 minutes to separate the beads from the supernatant. 200 µL supernatant was transferred to a new tube using a gel-loading tip, and the beads were washed with 100 µL TEAB buffer. The beads were centrifuged at 2000 rpm for 5 minutes and the supernatant combined with the previous. 120 µL acetonitrile was added to each supernatant sample before addition of 80 µL (200 µg) of TMT 10 plex (Thermo Scientific, cat #90110) reconstituted in acetonitrile. The samples were incubated at room temperature for 1 hour and vortexed occasionally. 7 µL of freshly prepared 5% hydroxylamine in water was added to each sample to quench the reaction, vortexed, and incubated for 15 minutes before quenching with addition of 5 µL MS-grade formic acid. The samples were then vacuum centrifuged to dryness. The samples were combined by redissolving the contents of one tube in 200 µL 0.1 % trifluoroacetic acid solution in water and sequential transfer to the respective multiplexed experiment until all samples were redissolved. This stepwise process was repeated with an additional 100 µL 0.1 % TFA solution for a final volume of 300 µL. The pooled samples were fractionated using the Pierce high pH Reversed-Phase Fractionation Kit (Thermo Fisher Scientific 84868) according to manufacturer’s instructions. The peptide fractions were eluted from the spin column with solutions of 0.1% triethylamine containing an increasing concentration of MeCN (5 - 95% MeCN; 8 fractions). Samples were dried via vacuum centrifugation, reconstituted in 50 µL 0.1% formic acid, and stored at -80 °C until ready for mass spectrometry analysis.

### Affinity precipitation and quantitative TMT-MuDPIT analysis of proteins covalently modified by AA147^yne^ and AA132^yne^

HEK293T-Rex or HepG2 cells in 10 cm plates were treated for 6 h with vehicle (0.1% DMSO), AA147^yne^ (10 μM), or AA132^yne^ (10 μM), at 37 °C. Lysates were prepared in radioimmunoprecipitation assay (RIPA) buffer (150 mM NaCl, 50 mM Tris pH 7.5, 1% Triton X-100, 0.5% sodium deoxycholate, and 0.1% SDS) with fresh protease inhibitor cocktail (Roche, Indianapolis, IN) and centrifuged for 20 min at 10000×g. Protein concentrations of supernatants were determined by the BCA assay (Thermo Fisher). For each sample, 100 μg of lysate were reacted with click reagents to give final concentrations as follows: 100 µM of diazo biotin-azide (Click Chemistry Tools, Scottsdale, AZ), 800 µM copper (II) sulfate, 1.6 mM BTTAA ligand (2-(4-((bis((1-tert-butyl-1H-1,2,3-triazol-4-yl)methyl)amino)methyl)-1H-1,2,3-triazol-1-yl)acetic acid) (Albert Einstein College), and 5 mM sodium ascorbate. The reaction was placed on a shaker at 1000 rpm at 30 °C for 2 h. The proteins were then precipitated from the reaction mixture by adding an equal volume of 3:1 chloroform/methanol. The pellet was washed three times with 1:1 chloroform/methanol. The precipitate was suspended in 500 µL of 6 M urea with 25 mM ammonium bicarbonate and 140 µL of 10% SDS was added to this mixture to help solubilize the protein. 50 µL of high-capacity streptavidin beads were washed with PBS and mixed with the protein solution in 6 mL of phosphate-buffered saline (PBS). This suspension was placed on a rotator or a shaker and agitated for 2 h. The beads were centrifuged and washed 5 times with PBS and 1% SDS. The protein was eluted from the beads by two washes of 50 mM sodium dithionite in 1% SDS for 1 h and then precipitated by chloroform/methanol precipitation as described above.

Air-dried pellets from the affinity precipitation were resuspended in 1% RapiGest SF (Waters) in 100 mM HEPES (pH 8.0). Proteins were reduced with 5 mM tris(2-carboxyethyl)phosphine hydrochloride (Thermo Fisher) for 30 min and alkylated with 10 mM iodoacetamide (Sigma Aldrich, St. Louis, MO) for 30 min at ambient temperature and protected from light. Proteins were digested for 18 h at 37 °C with 2 μg trypsin (Promega). After digestion, 20 μg of peptides from each sample were reacted for 1 h with the appropriate TMT-NHS isobaric reagent (ThermoFisher) in 40% (v/v) anhydrous acetonitrile and quenched with 0.4% NH4HCO3 for 1 h. Samples with different TMT labels were pooled and acidified with 5% formic acid. Acetonitrile was evaporated on a SpeedVac and debris was removed by centrifugation for 30 min at 18,000×g. MuDPIT (Multi-Dimensional Protein Identification Technology) microcolumns were prepared as described previously 79. LCMS/MS analysis was performed using a Q Exactive mass spectrometer equipped with an EASY nLC 1000 (Thermo Fisher). MuDPIT experiments were performed by 5 min sequential injections of 0, 20, 50, 80, 100% buffer C (500 mM ammonium acetate in buffer A) and a final step of 90% buffer C / 10% buffer B (20% water, 80% acetonitrile, 0.1% fomic acid, v/v/v) and each step followed by a gradient from buffer A (95% water, 5% acetonitrile, 0.1% formic acid) to buffer B. Electrospray ionization was performed directly from the analytical column by applying a voltage of 2.5 kV with an inlet capillary temperature of 275°C. Data-dependent acquisition of MS/MS spectra was performed with the following settings: eluted peptides were scanned from 400 to 1800 m/z with a resolution of 30,000 and the mass spectrometer in a data dependent acquisition mode. The top ten peaks for each full scan were fragmented by HCD using a normalized collision energy of 30%, a 100 ms activation time, a resolution of 7500, and scanned from 100 to 1800 m/z. Dynamic exclusion parameters were 1 repeat count, 30 ms repeat duration, 500 exclusion list size, 120 s exclusion duration, and exclusion width between 0.51 and 1.51. Peptide identification and protein quantification was performed using the Integrated Proteomics Pipeline Suite (IP2, Integrated Proteomics Applications, Inc., San Diego, CA) as described previously.

### RNA-seq analysis

Cells were lysed and total RNA collected using the Quick-RNA Miniprep kit from Zymo Research (R1055) according to manufacturer’s instructions. RNA concentration was then quantified by NanoDrop. Whole transcriptome RNA was then prepared and sequenced by BGI Americas on the BGI Proprietary platform, which provided paired-end 50 bp reads at 20 million reads per sample. Each condition was performed in triplicate. RNAseq reads were aligned using DNAstar Lasergene SeqManPro to the Homo_sapiens-GRCh38.p7 human genome reference assembly, and assembly data were imported into ArrayStar 12.2 with QSeq (DNAStar Inc.) to quantify the gene expression levels and normalization to reads per kilobase per million. Differential expression analysis was assessed using DESeq2 in R, which also calculated statistical significance calculations of treated cells compared to vehicle-treated cells using a standard negative binomial fit of the reads per kilobase per million data to generate fold-change quantifications. (doi: 10.1186/s13059-014-0550-8) Gene Ontology (GO) analysis was performed using Panther (geneontology.org).^56, 57^

### Statistical Analysis

Unless otherwise noted, the data were tested for significance using One-Way ANOVA with a post-hoc Dunnett’s test.

### General Synthetic Procedures

All compounds and reagents were purchased from Sigma-Aldrich, Acros, Alfa Aesar, Combi-blocks, and EMD Millipore unless otherwise noted and were used without further purification. Thin layer chromatography with Merck silica plates (60-F254), using UV light as the visualizing agent, was used to monitor reaction progress. Flash column chromatography was carried out using a Teledyne Isco Combiflash Nextgen 300+ machine using Luknova SuperSep columns (SiO2,25 µm) with ethyl acetate and hexanes as eluents. ^1^H NMR spectra were recorded on a Varian INOVA-400 400MHz spectrometer. Chemical shifts are reported in δ units (ppm) relative to residual solvent peak. Coupling constants (*J*) are reported in hertz (Hz). Characterization data are reported as follows: chemical shift, multiplicity (s=singlet, d=doublet, t=triplet, q=quartet, br=broad, m=multiplet), coupling constants, number of protons, mass to charge ratio. The compound’s identity was confirmed via high-resolution mass spectrometry.

### Synthesis of AA132

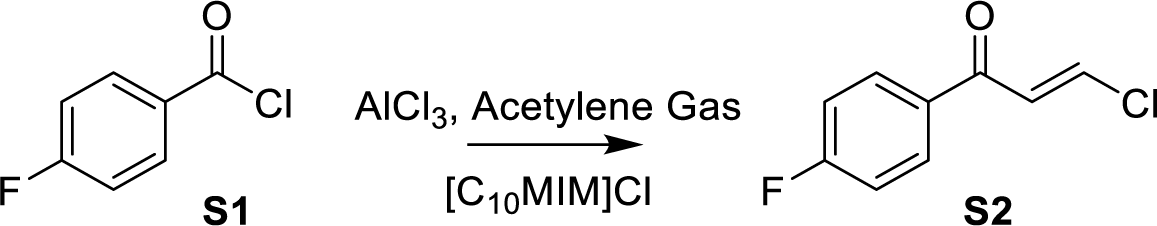

To a flame dried flask added 1-Decyl-3-methylimidazolium chloride (3.52 g, 13.6 mmol, 3.2 eq) and aluminum chloride (1.02 g, 7.65 mmol, 1.8 eq) and stirred for 16 hours under Argon to make the chloroaluminate fluid. To this flask at 0 °C, added 4-fluorobenzoyl chloride (S1, 500 µL, 4.25 mmol, 1 eq) slowly and the mixture became syrupy. The mixture was warmed to 75 ° C and acetylene gas generated in a separate flask from calcium carbide was bubbled through for 2 hours to afford the ß-chlorovinyl ketone. The crude was added to ice water and extracted with ether which was washed once with brine and concentrated to give a yellow oil (S2), which was used for the next step without further purification.

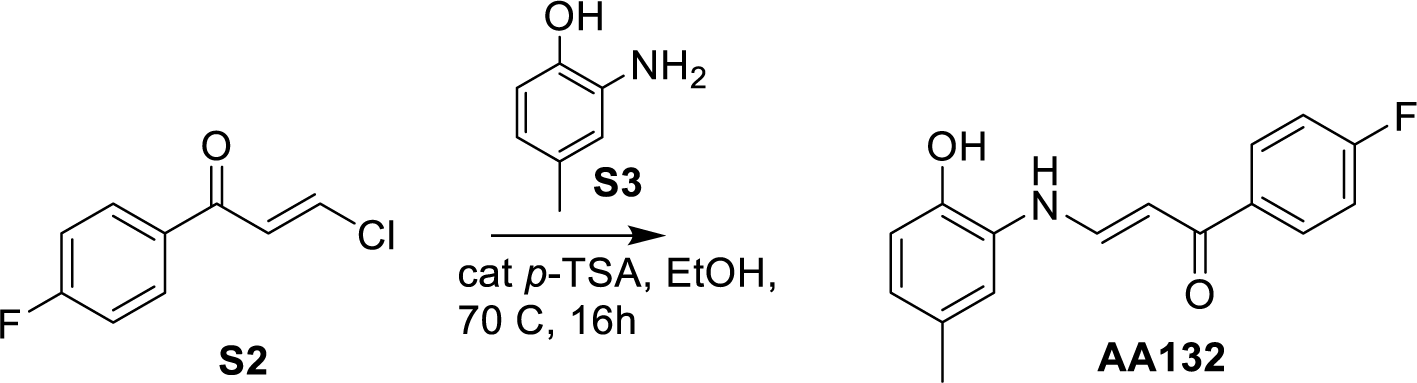

To a microwave vial with S2 (141 mg, 0.75 mmol, 1 eq), we added 2-amino-*p*-cresol (S3, 91 mg, 0.75 mmol, 1 eq) and *p*-toluenesulfonic acid (28.5 mg, 0.15 mmol,0.2 eq). Solids were dissolved in 3 mL EtOH and stirred under argon for 16h at 70 °C. Reaction diluted in EtOAc and washed sequentially with water and 1M HCl before drying over MgSO4. Product purified by column chromatography (SiO2, 4:1 Hex:EtOAc) to give AA132 as a yellow solid (98.2 mg, 48% yield).

^1^H NMR (400 MHz, Acetone) δ 12.28 (d, J = 12.8 Hz, 1H), 8.82 (d, J = 0.6 Hz, 1H), 8.13 – 8.01 (m, 2H), 7.86 (ddd, J = 12.8, 7.8, 0.5 Hz, 1H), 7.29 – 7.19 (m, 3H), 6.88 (d, J = 8.1 Hz, 1H), 6.74 (dtd, J = 8.0, 1.4, 0.7 Hz, 1H), 6.13 (d, J = 7.8 Hz, 1H), 2.28 (d, J = 0.7 Hz, 3H).

^19^F NMR (376 MHz, Acetone) δ -113.86 ^13^C NMR (100 MHz, ACETONE-D6) δ 188.04, 165.98, 163.50, 144.29, 143.67, 136.15, 136.12, 129.89, 129.80, 129.78, 128.54, 123.92, 115.44, 115.28, 115.07, 114.47, 92.78, 20.07.

**HRMS:** calculated for C16H14FNO2[M+ H^+^] 272.1087, found 272.1085

### Synthesis of AA132^yne^

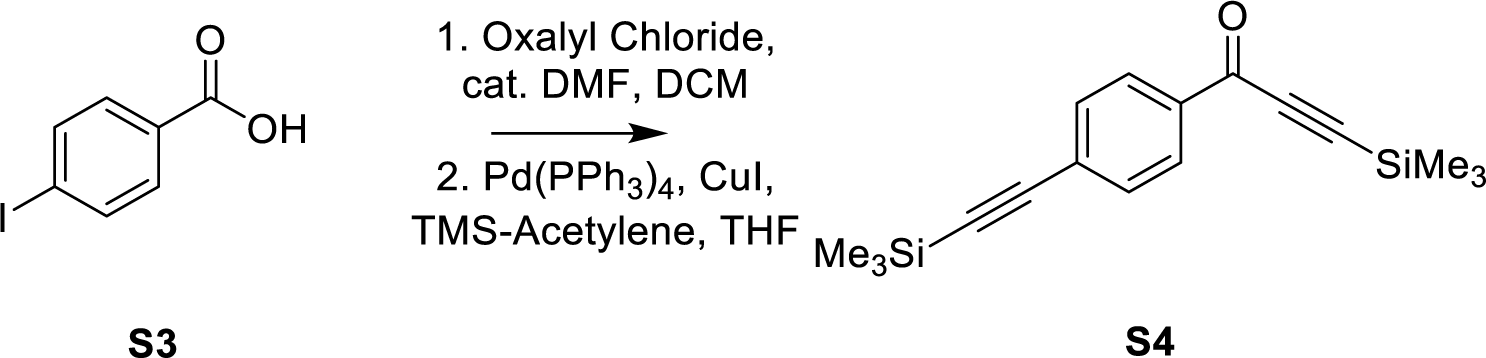

To 4-Iodobenzoic acid (S3, 738 mg, 3 mmol, 1 eq) dissolved in 15 mL DCM added oxalyl chloride (321 µL, 3.75 mmol, 1.25 eq) dropwise at 0 °C before addition of several drops of DMF. Reaction allowed to warm to room temperature and stir for 3h before TLC indicated complete conversion to the acyl chloride. Solvent removed under reduced pressure to give a yellow-white powder. To flask containing crude residue added copper iodide (50 mg, .11 mol%) and Palladium-tetrakis(triphenylphosphine) (80 mg, 2 mol %), then dissolved in 10 mL anhydrous THF. After 3x freeze-pump-thaw cycles, added diisopropylethylamine (2.6 mL, 15 mmol, 5 eq), and trimethylsilylacetylene (980 µL, 3.3 mmol, 1.1 eq). Reaction stirred under argon at 40 °C overnight before washing with saturated NaHCO3, and brine before drying over MgSO4 and concentrating. The crude residue was purified by flash column chromatography (SiO2, 9:1 Hex/EtOAc) to give an orange solid (S4, 638 mg, 71 % yield).

^1^H NMR (400 MHz, Acetone) δ 8.18 – 8.08 (m, 2H), 7.70 – 7.61 (m, 2H), 0.35 (d, J = 1.1 Hz, 9H), 0.28 (d, J = 1.0 Hz, 9H).

^13^C NMR (100 MHz, Acetone-D6) δ 175.96, 136.06, 132.20, 129.41, 128.95, 103.96, 100.51, 98.61, - 1.02, -1.49.

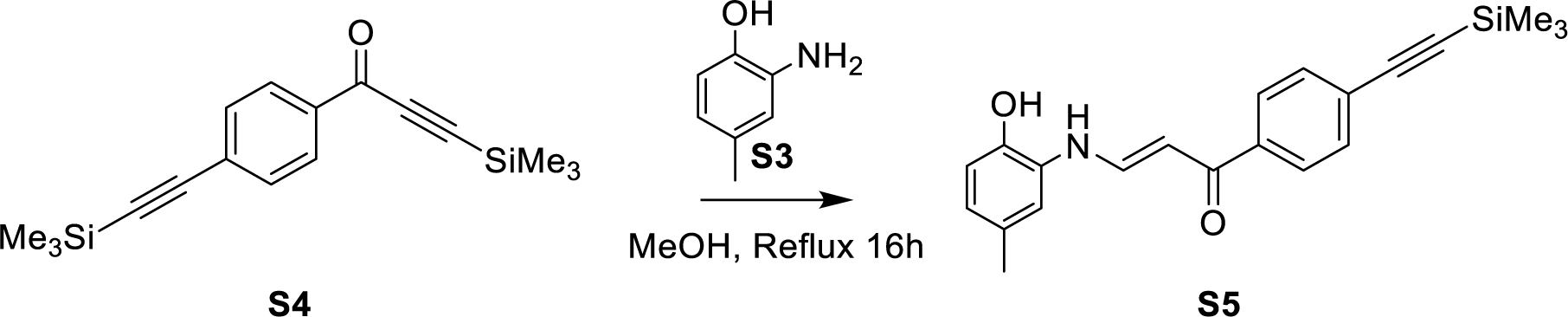

To a microwave vial charged with S4 (59 mg, 0.2 mmol, 1 eq), we added 2-amino-*p*-cresol (S3; 23.6 mg, 0.2 mmol, 1 eq). The reaction vessel was sealed before addition of 1 mL anhydrous methanol and heated at 70 ° C for 16 h. Reaction diluted in EtOAc, washed with 1M HCl and brine, and dried over MgSO4. The solvent was removed under reduced pressure. The crude residue purified by flash column chromatography (SiO2, 9:1 Hex/EtOAc) to afford the product S5 as a yellow powder (43 mg, 62% yield).

^1^H NMR (500 MHz, Acetone) δ 12.36 (d, *J* = 12.8 Hz, 1H), 8.86 (s, 1H), 8.03 – 7.97 (m, 2H), 7.93 – 7.85 (m, 1H), 7.58 – 7.54 (m, 2H), 7.25 (d, *J* = 1.8 Hz, 1H), 6.88 (d, *J* = 8.1 Hz, 1H), 6.75 (ddd, *J* = 8.1, 2.0, 0.8 Hz, 1H), 6.16 (d, *J* = 7.8 Hz, 1H), 2.28 (d, *J* = 0.7 Hz, 3H), 0.27 (s, 9H).

^13^C NMR (126 MHz, Acetone) δ 205.25, 188.20, 144.44, 143.67, 139.40, 131.72, 129.73, 128.42, 127.27, 127.25, 125.74, 123.99, 115.42, 114.52, 104.58, 96.07, 93.00, 28.79, 28.64, 28.48, 19.99, - 1.02.

**HRMS:** calculated for C17H22OSi2 [M+ H^+^] 299.1287, found 299.1289

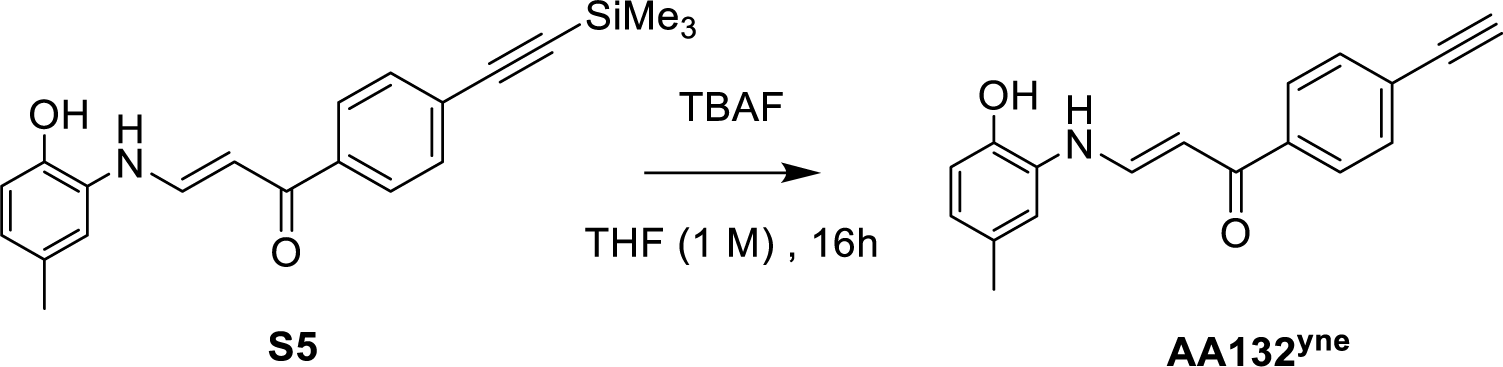

To S5 (0.1 mmol, 35 mg) dissolved in 0.5 mL THF, we added 0.5 mL of tetrabutylammonium fluoride (1 M) in THF overnight dropwise. Reaction was allowed to stir overnight at room temperature. Reaction diluted in EtOAc and washed with 1M HCl. The organic layer was collected, dried over MgSO4, and solvent removed under reduced pressure. The crude was purified by flash column chromatography (SiO2 4:1 Hex: EtOAc) to afford the product AA132^yne^ as a yellow powder (23.3 mg, 86 % yield).

^1^H NMR (400 MHz, ACETONE-*D*6) δ 12.32 (d, *J* = 12.9 Hz, 1H), 8.89 (s, 1H), 8.01 – 7.90 (m, 2H), 7.85 (dd, *J* = 12.8, 7.8 Hz, 1H), 7.59 – 7.53 (m, 2H), 7.21 (d, *J* = 1.9 Hz, 1H), 6.84 (d, *J* = 8.1 Hz, 1H), 6.76– 6.67 (m, 1H), 6.12 (d, *J* = 7.7 Hz, 1H), 3.81 (s, 1H), 2.23 (s, 3H).

^13^C NMR (100 MHz, ACETONE-*D*6) δ 205.44, 188.31, 144.55, 143.75, 139.62, 132.00, 129.77, 128.44, 127.37, 125.09, 124.09, 115.47, 114.56, 93.05, 82.99, 80.54, 20.08.

**HRMS:** calculated for C18H15NO2 [M+ H^+^] 278.1181, found 278.1187

### NMR Spectra for AA132

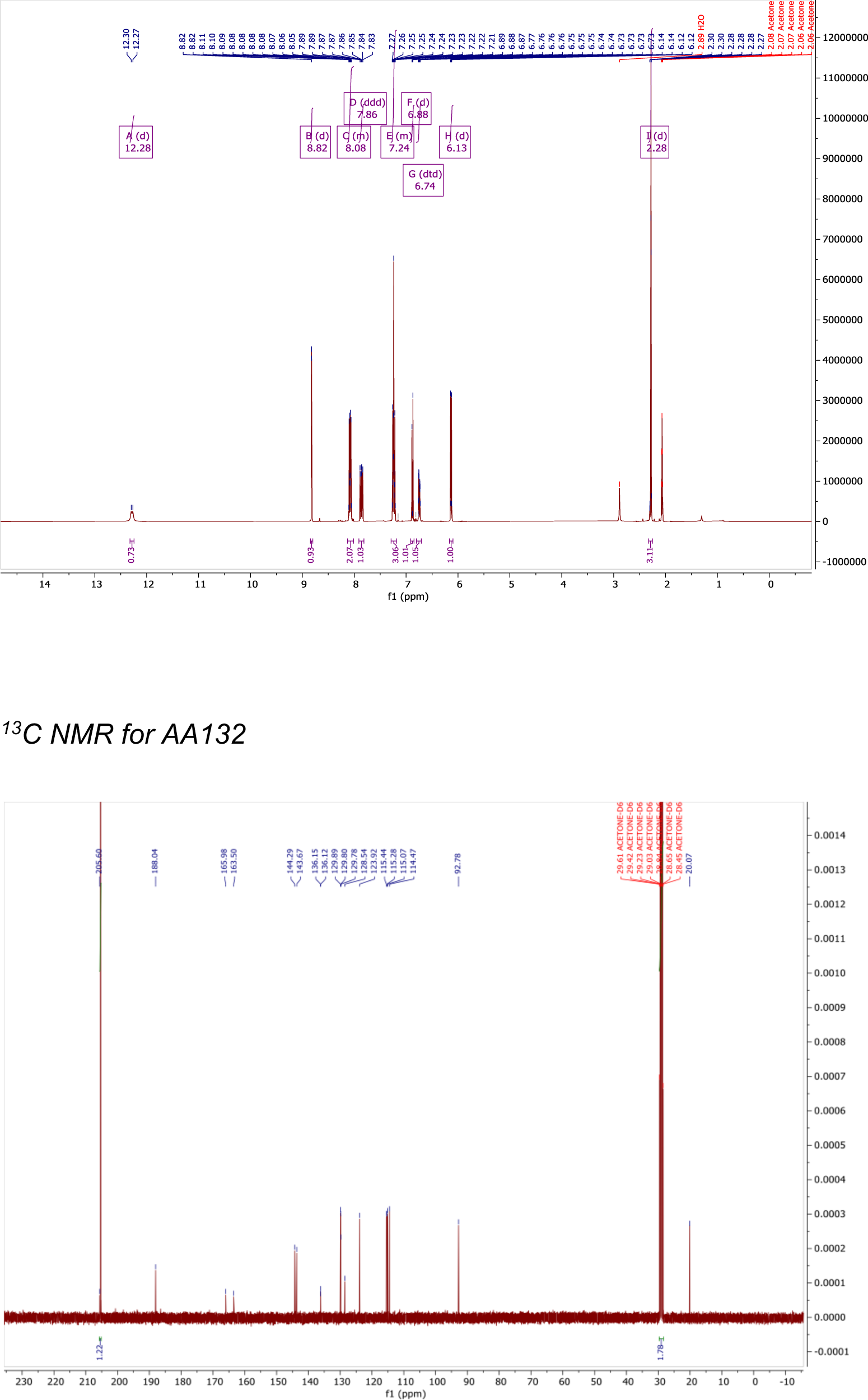
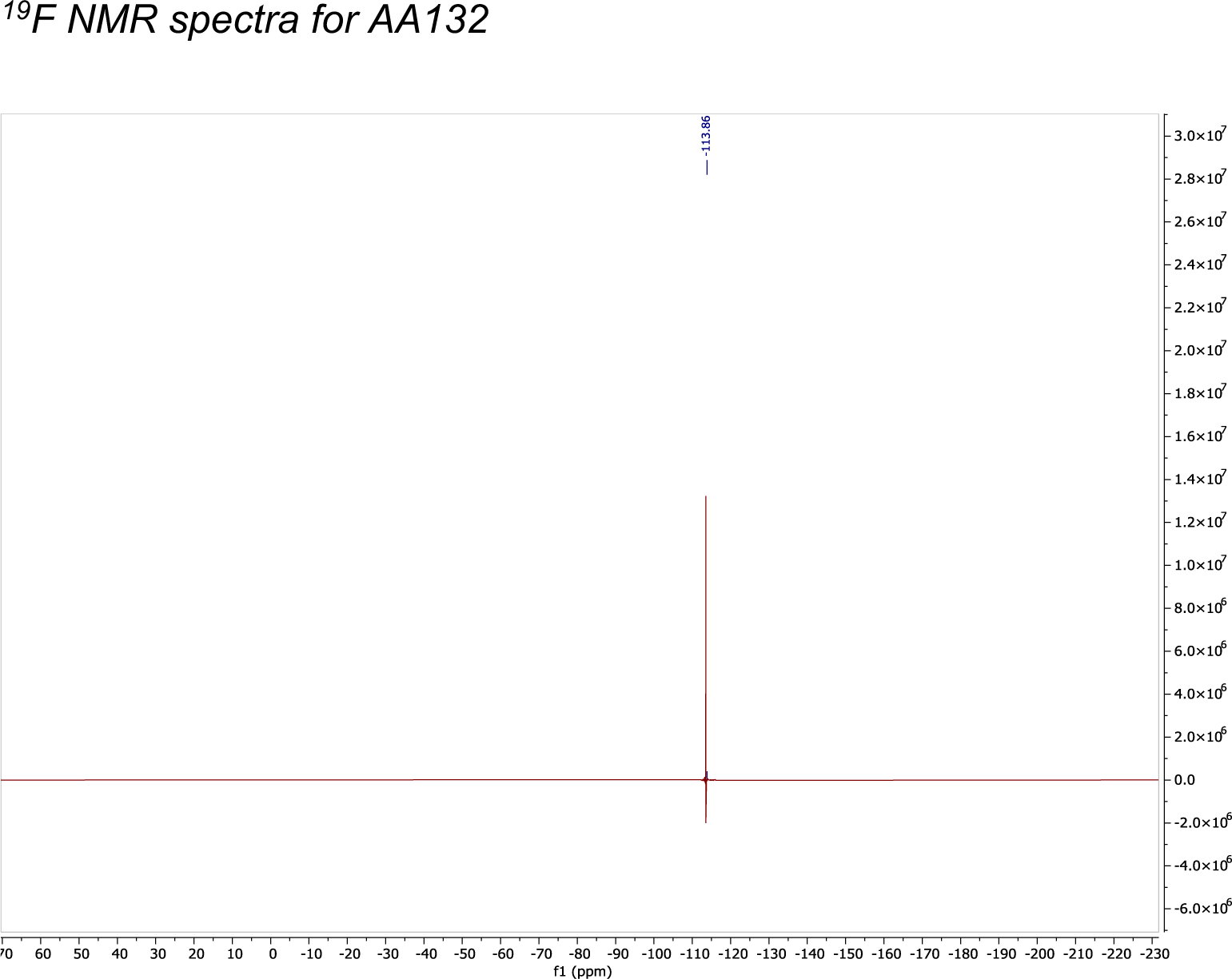

### NMR Spectra for S5

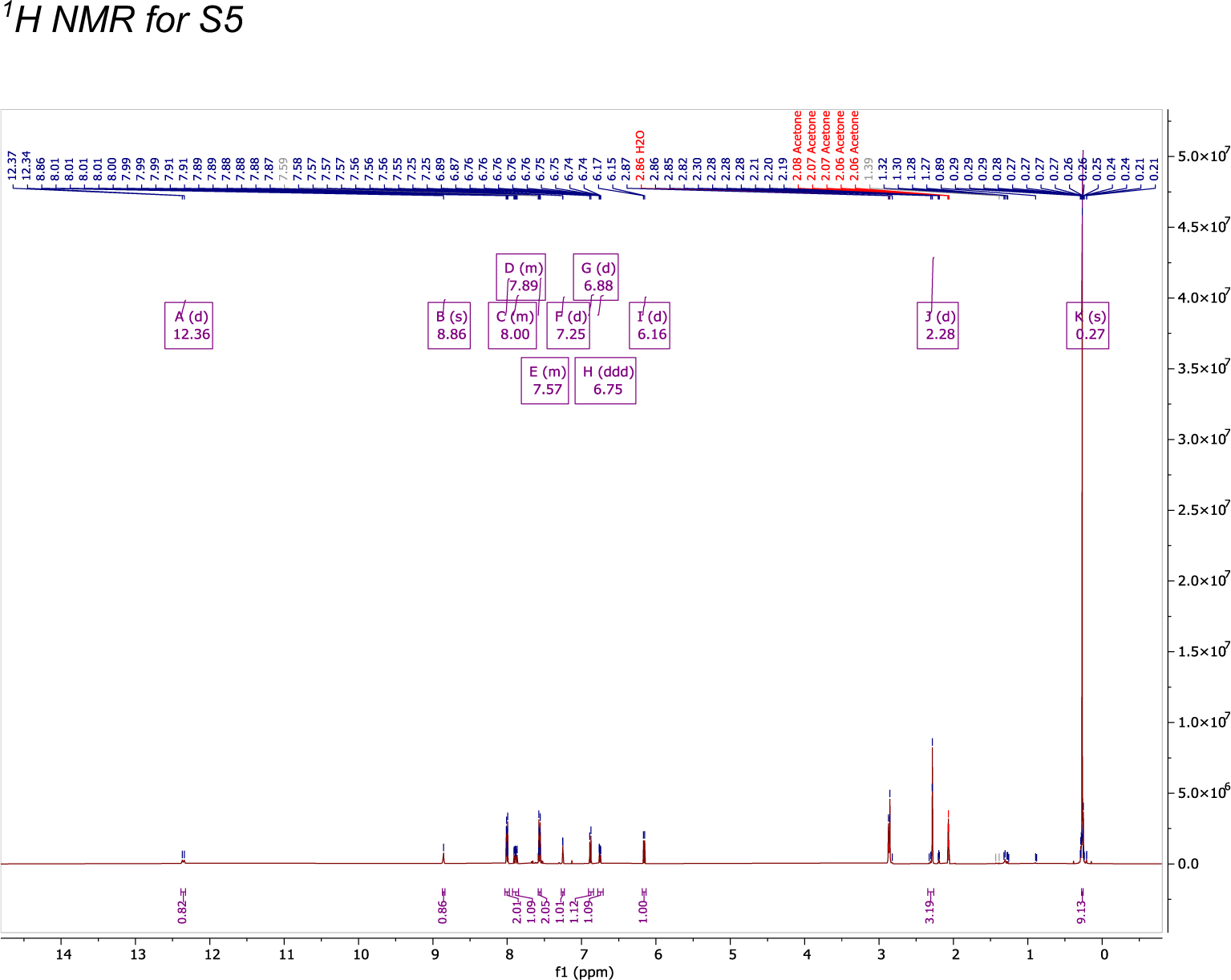
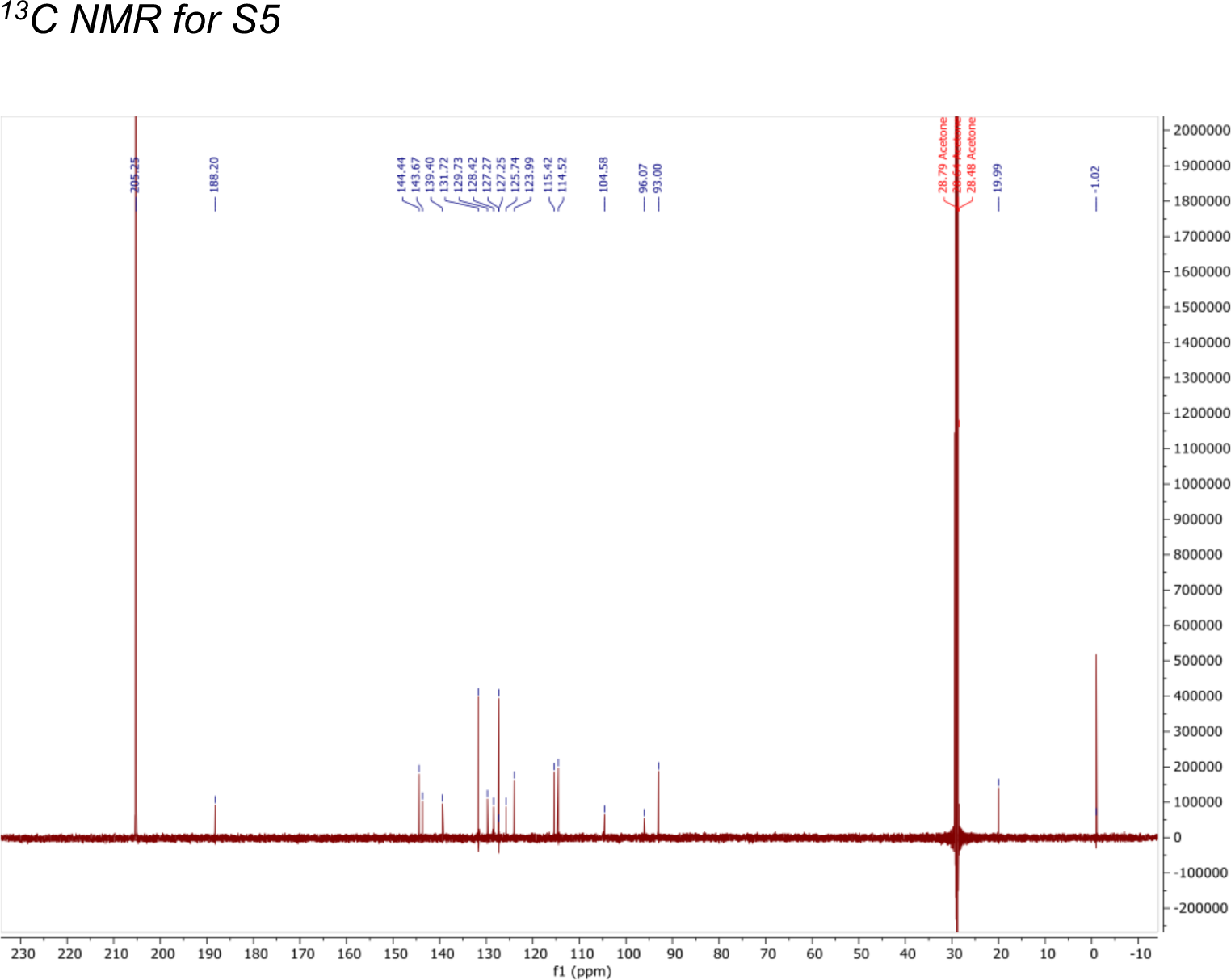

### NMR Spectra for AA132^yne^

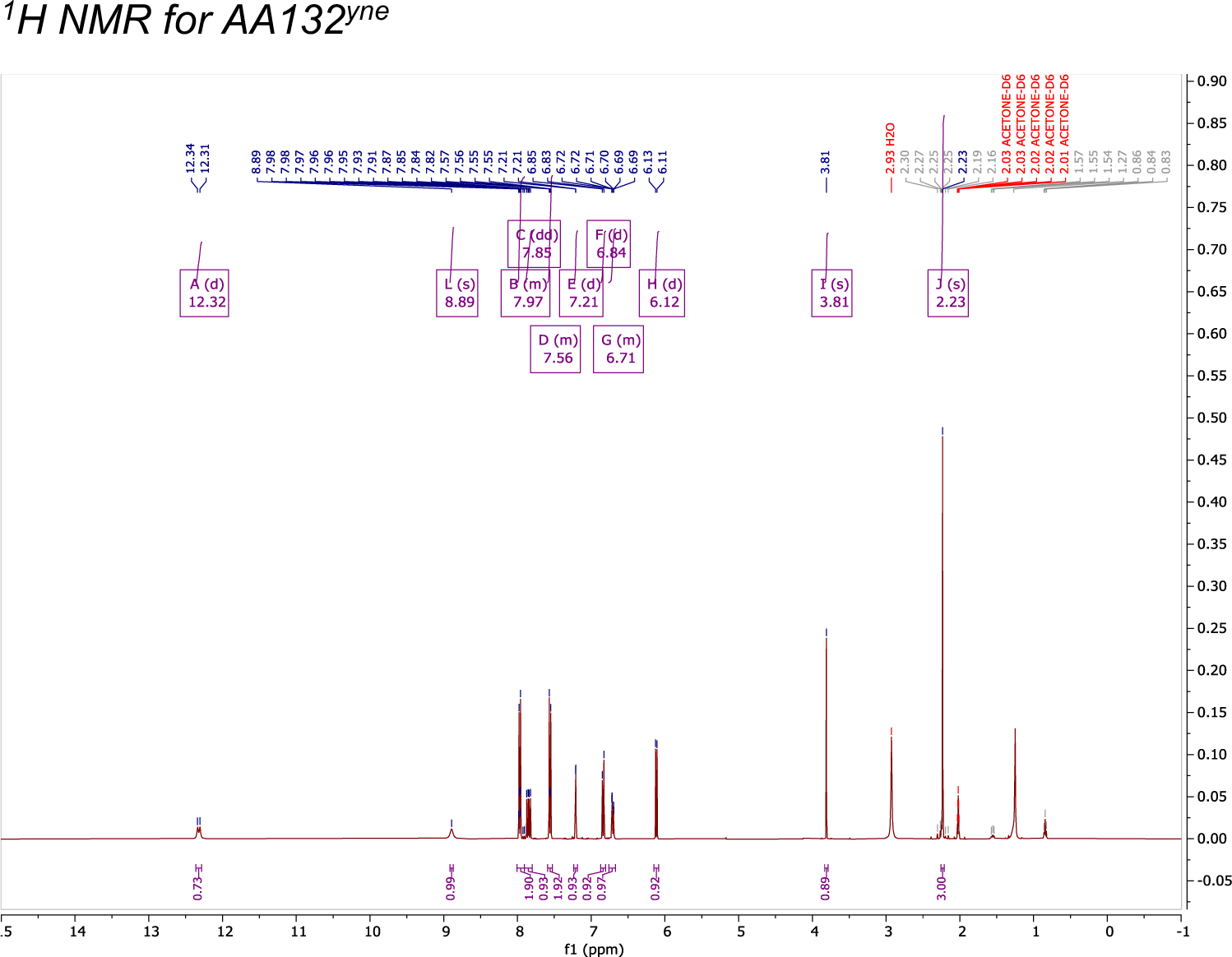
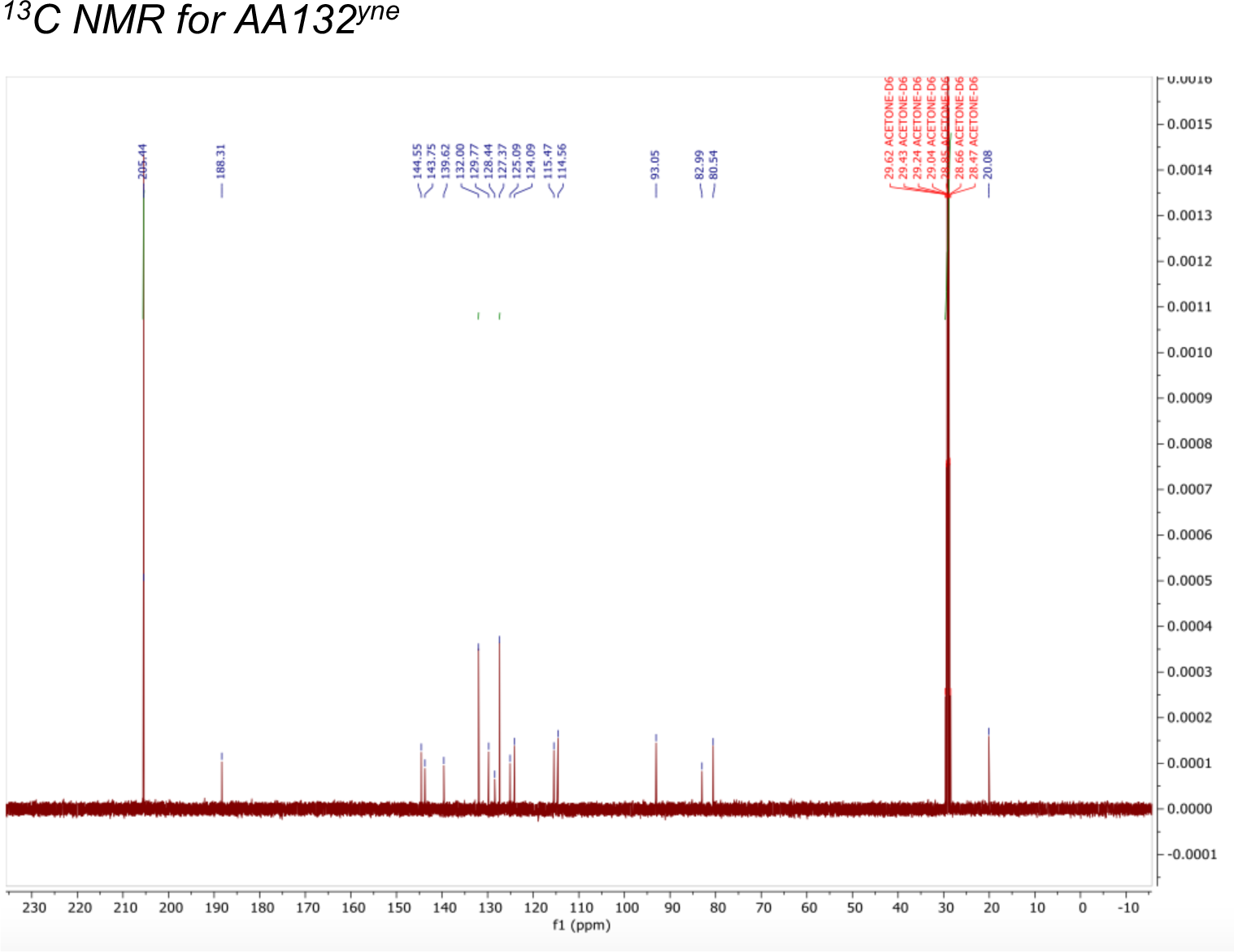

**Supplementary Figure 1.**
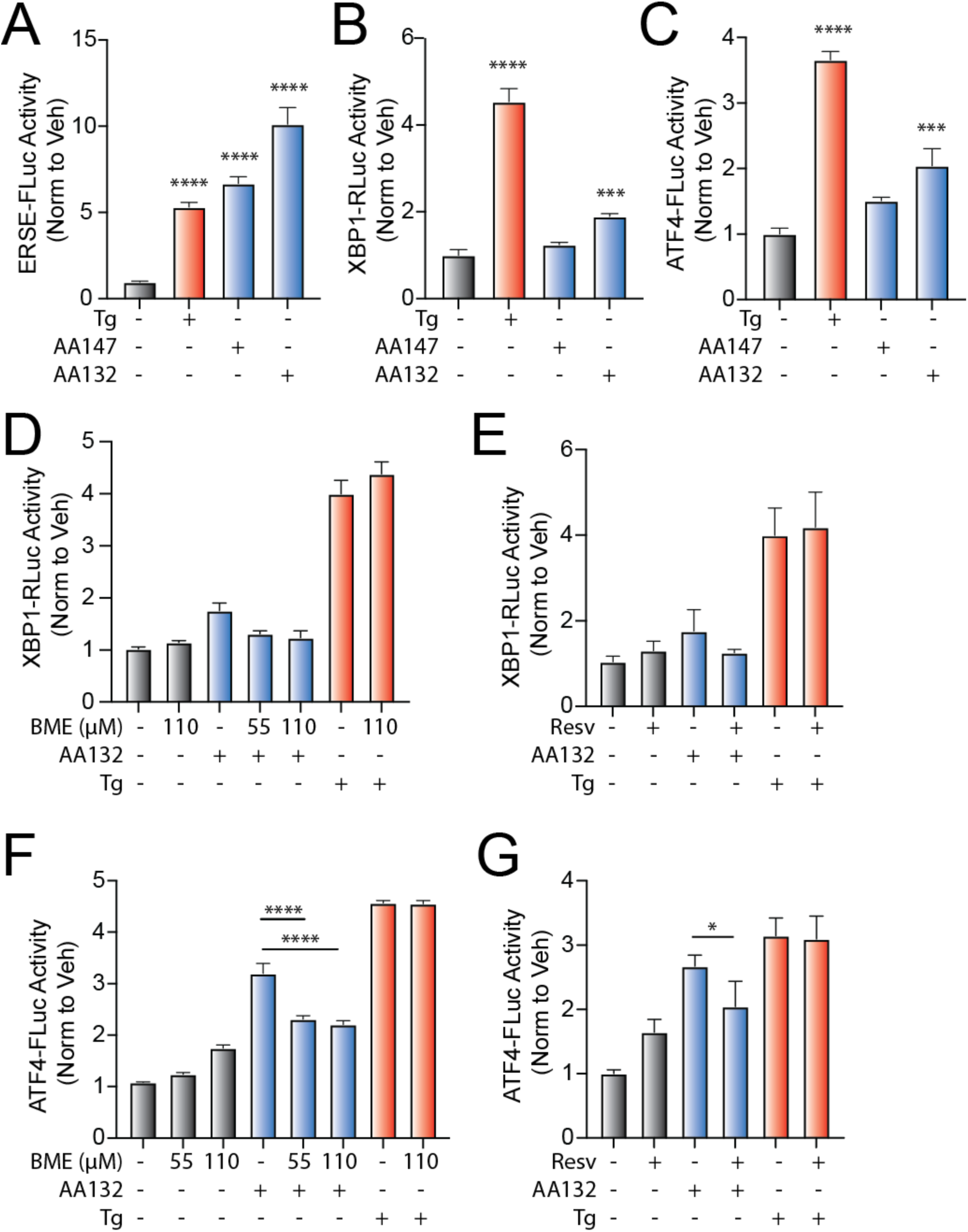
AA132 activates ATF6 signaling pathways through a mechanism involving metabolic activation and covalent protein modification. **A.** Bar graph showing the activation of the ERSE.FLuc ATF6 reporter in HEK293T cells treated with Veh (0.1% DMSO), thapsigargin (Tg; 500 nM), AA147 (10 µM), or AA132 (10 µM) for 18 hr. Error bars show SEM for 6 independent experiments. \*\*\*\**p* < 0.0001. **B.** Bar graph showing the activation of the XBP1s.RLuc IRE1 reporter in HEK293T cells treated with Veh (0.1% DMSO), Tg (500 nM), AA147 (10 µM), or AA132 (10 µM) for 18 hr. Error bars show SEM for 6 independent experiments. \*\*\**p* < 0.001, \*\*\*\**p* < 0.0001. **C.** Bar graph showing the activation of the ATF4.FLuc PERK reporter in HEK293T cells treated with Veh (0.1% DMSO), Tg (500 nM), AA147 (10 µM), or AA132 (10 µM) for 18 hr. Error bars show SEM for 6 independent experiments. \*\*\**p* < 0.001, \*\*\*\**p* < 0.0001. **D.** Bar graph showing the activation of the XBP1s.RLuc IRE1 reporter in HEK293T cells treated with AA132 (10 µM) or Tg (500 nM) in the presence or absence of β-mercaptoethanol (BME; 55 μM or 110 μM) for 18 hr. Error bars show SEM for 6 independent experiments**. E.** Bar graph showing the activation of the XBP1s.RLuc IRE1 reporter in HEK293T cells treated with AA132 (10 µM) or Tg (500 nM) in the presence or absence of resveratrol (2.5 µM) for 18 hr. Error bars show SEM for 6 independent experiments. **F.** Bar graph showing the activation of the ATF4.FLuc PERK reporter in HEK293T cells treated with AA132 (10 µM) or Tg (500 nM) in the presence or absence of β-mercaptoethanol (BME; 55 μM or 110 μM) for 18 hr. Error bars show SEM for 6 independent experiments. \*\*\*\**p* < 0.0001 **G.** Bar graph showing the activation of the ATF4.FLuc PERK reporter in HEK293T cells treated with AA132 (10 µM) or Tg (500 nM) in the presence or absence of resveratrol (2.5 µM) for 18 hr. Error bars show SEM for 6 independent experiments. \**p* < 0.05.

**Supplementary Figure 2.**
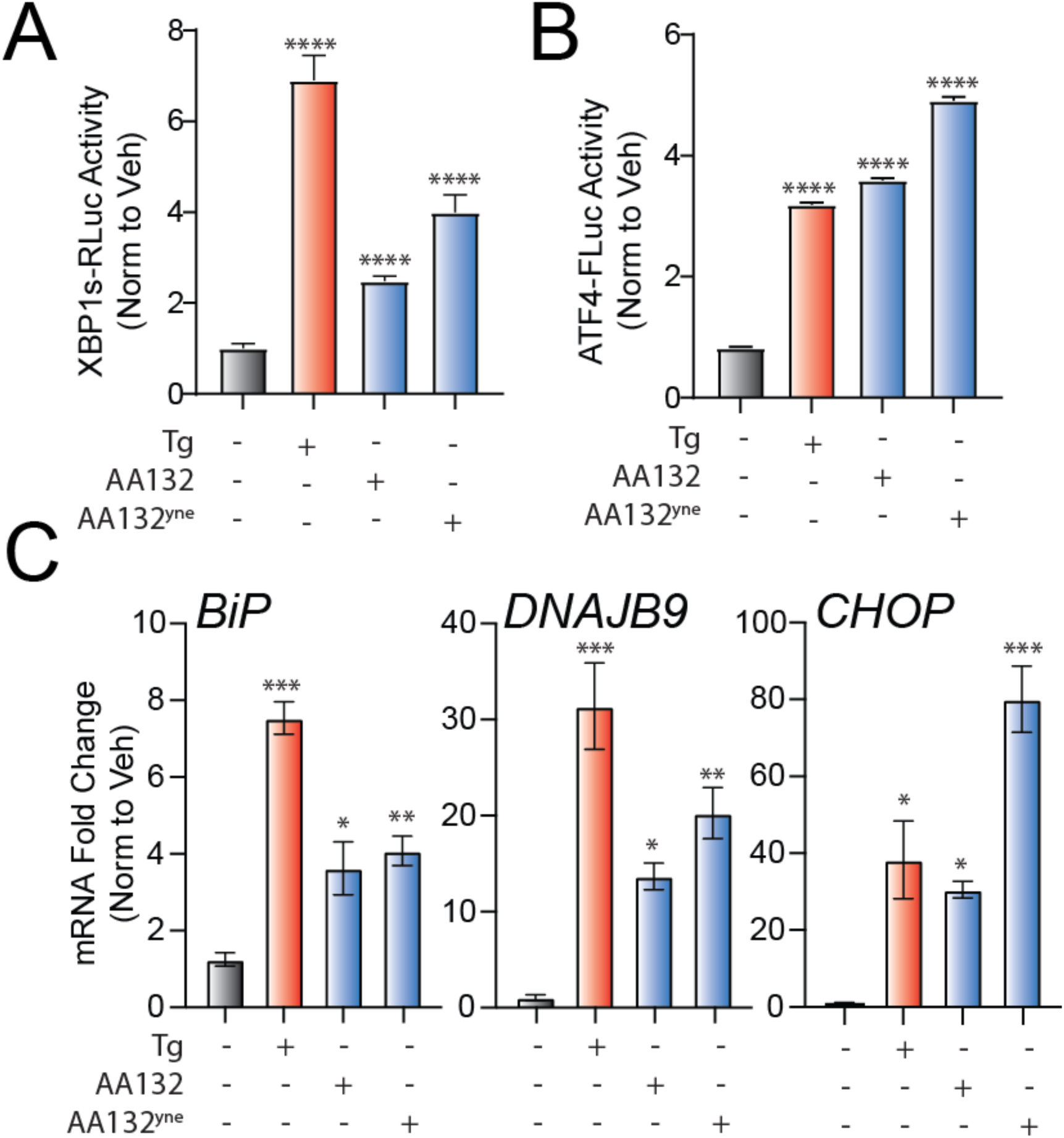
Development of a Functional Affinity Enrichment Probe for AA132. **A.** Bar graph showing the activation of the XBP1s.RLuc IRE1 reporter in HEK293T cells treated with Veh (0.1% DMSO), thapsigargin (Tg; 500 nM), AA132 (10 µM), or AA132^yne^ (10 µM) for 18 hr. ****p<0.0001 **B.** Bar graph showing the activation of the ATF4.FLuc reporter in HEK293T cells treated with Veh (0.1% DMSO), Tg (500 nM), AA132(10 µM), or AA132^yne^ (10 µM) for 18 hr ****p<0.0001 **C.** Graph showing qPCR of the ATF6 target gene *BiP*, PERK target gene *CHOP*, and XBP1s target gene *DNAJB9* in MEF cells treated for 6 h with the indicated compound (10 µM). N = 3 biological replicates. *p<0.05, **p<0.01, ***p<0.001.

**Supplementary Figure 3.**
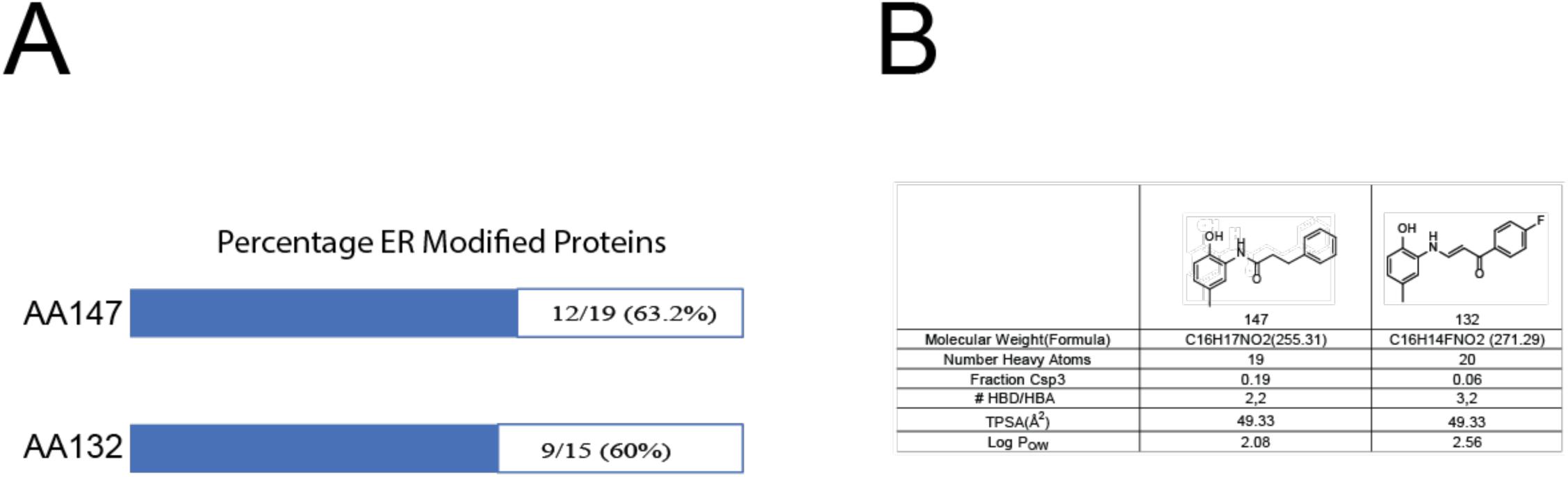
AA132^yne^ Covalently Modifies ER PDIs. **A.** Graph showing proportion of AA147^yne^ and AA132^yne^ target proteins localized to the ER. **B.** Calculated physicochemical properties of AA147 and AA132. Values calculated using SwissADME (SwissADME.ch). TPSA = Total Polar Surface Area; HBD = Hydrogen Bond Donors; HBA = Hydrogen Bond Acceptors.

**Supplementary Figure 4.**
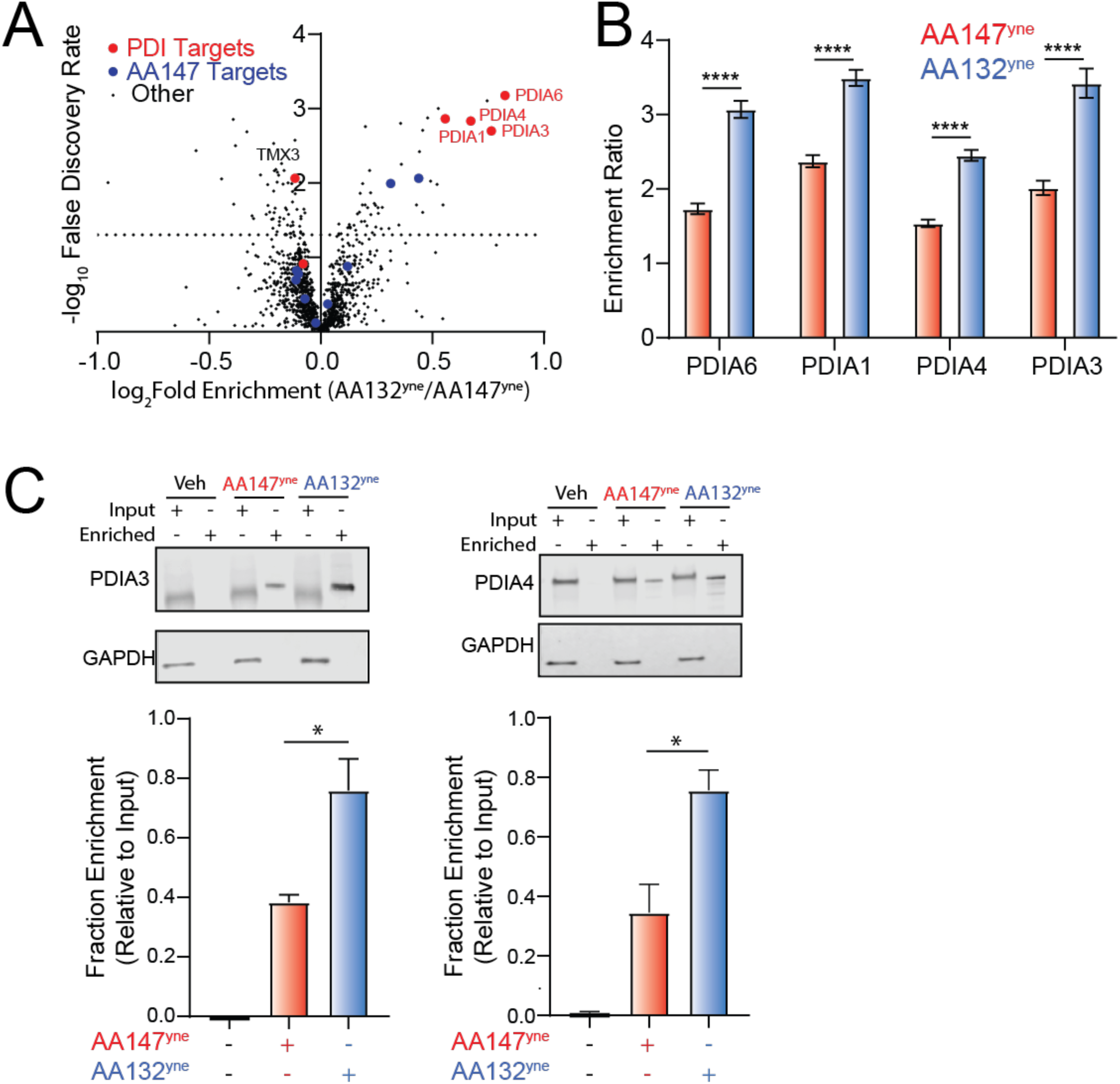
AA132^yne^ Shows Higher PDI Labeling as Compared to AA147^yne^. **A.** Volcano plot showing log2 fold enrichment of AA132^yne^ labeled proteins relative to AA147^yne^ labeled proteins (x-axis) versus the –log FDR (y-axis) in HepG2 cells (10 µM, 6h). Proteins with GO annotation for PDI (GO: 0003756) labeled in red and additional previously defined AA147^yne^ targets labeled in blue. Data shown in **Table S2**. **B.** Bar graph of enrichment ratio of select PDIs by indicated the compound relative to DMSO from data shown in **Fig. S4A** (N = 4 biological replicates). ****p < 0.001 from multiple unpaired t-test. **C**. Representative immunoblot and quantification of PDIA3 and PDIA4 recovery in streptavidin enrichments from HEK293T cells treated with AA147^yne^ (10 µM; 6 h) or AA132^yne^ (10 µM; 6 h) and then conjugated to biotin. Fraction enrichment was calculated by dividing the signal in enriched samples by the input signal. *p < 0.05 from unpaired t test for N=3 replicates.

**Supplementary Figure 5.**
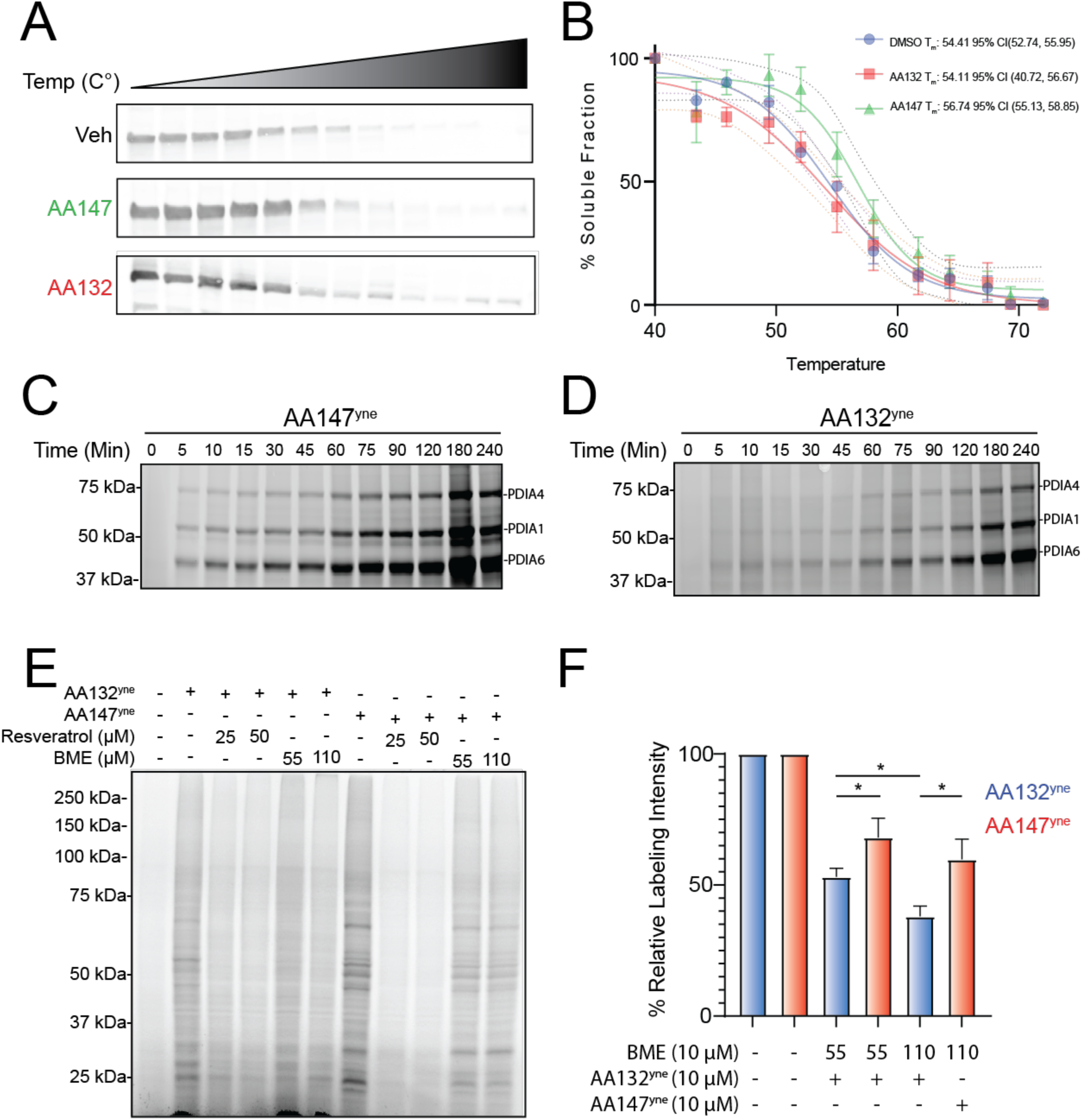
AA132^yne^ Shows Slower Protein Labeling Kinetics as Compared to AA147^yne^. **A.** Representative immunoblot of the soluble fraction of PDIA1 from heat-treated ALMC2 cells at the temperatures indicated (40-72°C) preincubated with listed compound (10 µM, 2h). **B.** Graph of percent soluble fraction (y-axis) versus temperature (x-axis) for data in **Fig S5A**. Fitted curves calculated using Boltzmann Sigmoidal Fit in Prism and plotted with 95% confidence intervals (dotted lines). Tm is temperature on calculated sigmoidal curve with 50% soluble PDIA1 fraction remaining. Error bars represent S.E.M (N = 3 biological replicates). **C.** Representative SDS-PAGE gel of Cy5-conjugated proteins from ALMC2 cells treated at indicated time point with AA147^yne^ (10 µM). **D.** Representative SDS-PAGE gel of Cy5-conjugated proteins from ALMC2 cells treated at indicated time point with AA132^yne^ (10 µM). **E.** Representative gel of AA132^yne^ and AA147^yne^ labeled proteins in HEK293T cells cotreated with indicated concentrations of β-mercaptoethanol or resveratrol for 4h. **F.** Quantification of **Fig S5E**. Error bars represent standard error of mean. *p* values calculated using two-sided Student T Test. Percent labeling calculated as lane intensity relative to cotreatment with vehicle (0.1% DPBS) \**p* < 0.05, **p < 0.01.

**Supplementary Figure 6.**
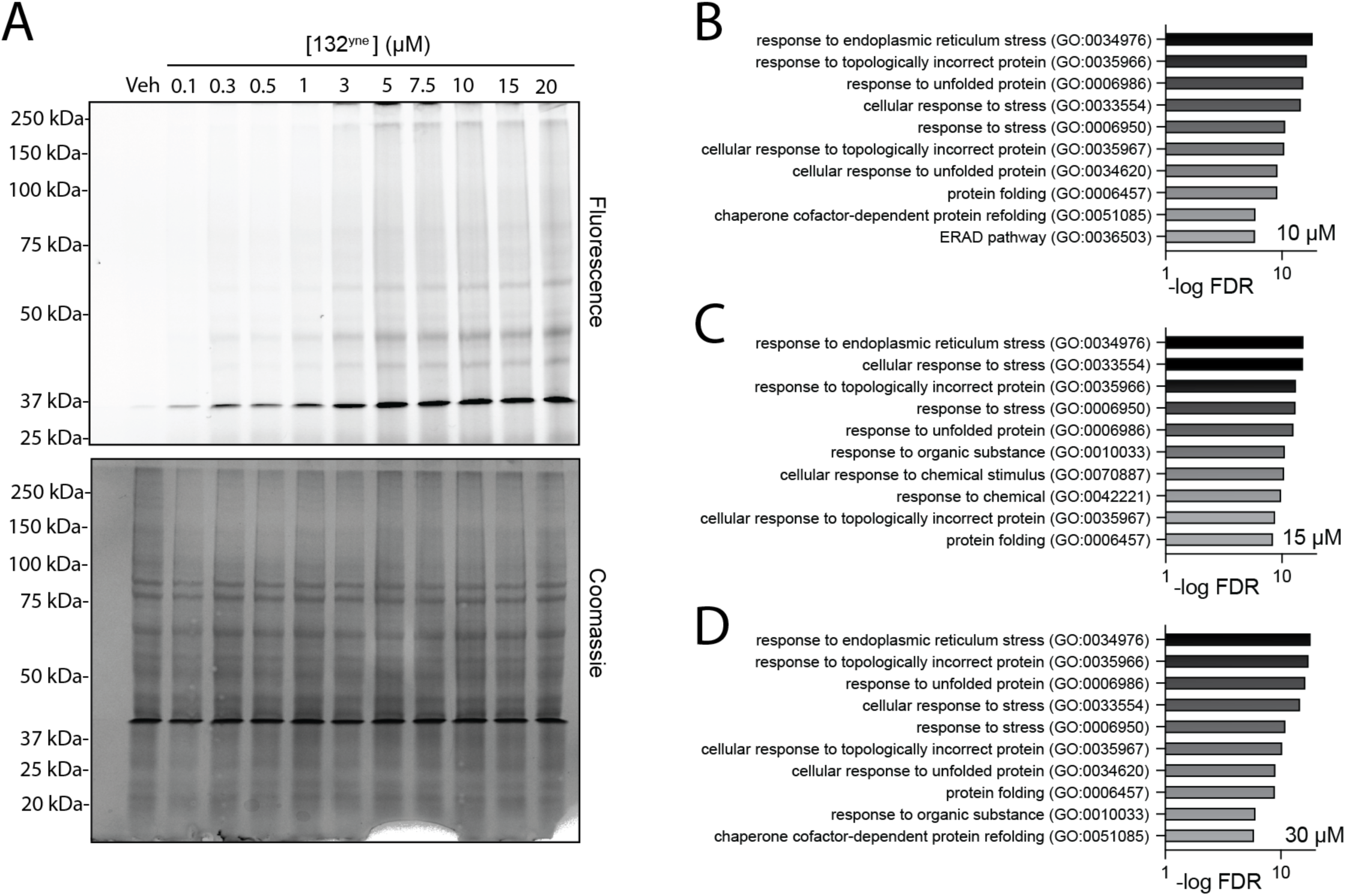
AA132 Selectively Activates ATF6 Transcriptional Signaling at Lower Doses. **A.** Fluorescence and coomassie-stained SDS-PAGE of lysates prepared from HEK293T cells treated with the indicated concentration of AA132^yne^ (4 h) and then conjugated to azide-cyanine. **B-D**. Top-10 GO terms for significantly induced genes (fold change >1.3, p<0.05) identified by RNAseq in HEK293T cells treated with 10 µM (**A**), 15 µM (**B**), or 30 µM (**C**) AA132 for 6 h. RNAseq data is included in **Table S3** Full GO analysis is included in **Table S4**.

